# ProFORGE: a composable sequence–structure–dynamics workflow validated by mechanoprotein design

**DOI:** 10.64898/2026.11.25.769315

**Authors:** Yanzhang Huang, Wanqing Yang, Yang Wang, Zilong Guo, Jie-Li Wang, Yanwei Wang, Hu Chen, Jianwei Shuai, Fangfu Ye

## Abstract

Designing a protein to resist tension requires identifying the contacts that bear the load. We developed ProFORGE to track residues as a scaffold changes length and to select designs by steered molecular dynamics. Starting with the 56-residue GB1 domain, we generated 14,542 candidates using protein language models for scaffold expansion and fixed-backbone design for sequence refinement. Of 182 sequences that passed the initial screen in the final round, 19 also met the AlphaFold 3 confidence threshold. The number of backbone hydrogen bonds between the terminal strands correlated with mean peak force at both simulated pulling speeds (Spearman ***ρ*** = 0.83–0.85). Two 98-residue designs shared less than 10% global sequence identity with wild-type GB1. In single-molecule magnetic-tweezers measurements, their most probable unfolding forces were 117.9 and 132.1 pN, or 5.5 and 5.8 times those of the wild-type domains measured on the same tethers.

---

Two protein domains in series experience the same tension, yet one can unfold while the other remains folded. For an elastomer, that difference determines when stretching releases additional chain length. For a molecular force probe, an unfolding step can report that a particular load range has been reached. Designing either response requires knowing which contacts bear the load and how readily they rupture when the protein is pulled from its attachment points. Multimodal protein language models can generate sequences from partial structural information ^1^, structure predictors can estimate the resulting folds ^2–4^, and inverse-folding networks can design sequences for a chosen backbone ^5^. These advances in modelling protein structure and fitness ^6,7^ have made it possible to connect generation, refinement and scoring in automated design workflows ^8–10^. A fold prediction, however, gives no direct measure of the force needed to unfold a domain under a specified loading protocol.

The B1 domain of streptococcal protein G (GB1) makes this distinction experimentally accessible. Its crystal structure (1PGA) ^11^ and NMR structure (3GB1) ^12^ place a four-stranded *β*-sheet against an *α*-helix. Pulling the N- and C-termini apart loads two parallel strands, *β*1 and *β*4, in shear across an interface spanned by 5–6 backbone hydrogen bonds ^13–17^. Changes in core packing or surface contacts can alter the force response of this 56-residue domain ^13–16^. Extending the interface poses a different design question. Can the two strands grow together while preserving their register and their packing against the rest of the fold? GB1 also provides its own experimental reference. Wild-type and designed domains can be placed in the same polyprotein and pulled under the same force calibration ^14,17^.

Several approaches already target protein mechanics directly. ForceGen conditions sequence generation on a desired unfolding response ^18^. Other studies have combined computation and experiment to strengthen helical scaffolds and titin domains ^19,20^, extended the force-bearing strands of titin I27 to increase hydrogen bonding ^21^, or redesigned the hydrophobic cores of helical bundles ^22^. Steered molecular dynamics (SMD) can trace how contacts rupture along a chosen pulling direction ^23^. For iterative backbone design, the challenge is to use that information in choosing the next scaffold. An insertion shifts the indices of the residues that define a structural constraint. A candidate can also retain a plausible overall fold while losing the strand contacts that motivated the edit. Across successive rounds, the design procedure therefore needs to keep track of the same interface and compare candidates under a common mechanical test.

We developed ProFORGE (Protein-language-model-guided Fold-aware Optimization, Refinement, and Generative Exploration) to couple these two operations. A residue map carries the structural anchors from parent to child as ESM-3 expands the scaffold, so strand continuity can be checked at the intended positions after each edit. Among candidates that pass structure and solubility screens, two stages of pulling in GROMACS ^24^ guide the choice of the parent for the next expansion. Once the backbone length is fixed, ProteinMPNN refines the sequence and a second structure predictor checks the final candidates. In this way the simulated response of one generation influences which backbone is edited in the next, while each change at the load-bearing interface can be traced to its parent. Starting from GB1, we followed that interface from 56 to 98 residues and used single-molecule magnetic tweezers to test whether the gain in simulated resistance persisted under slow experimental loading.

## Results

### The ProFORGE workflow and choice of sequence generators

ProFORGE represents a design round as a directed graph. Nodes import a structure, edit residue-level prompts, generate sequences and predict their folds. A user can connect these nodes to screening, fixed-backbone refinement and export, or add a custom scoring or analysis node through the same input and output ports (Fig. 1a,b and Supplementary Fig. 1a). In Protein Studio, the three-dimensional structure appears alongside residue-level tracks for sequence, coordinates, secondary structure, solvent-accessible surface area (SASA) and functional annotations (Supplementary Fig. 1b). After an insertion, deletion or splice, the software updates the correspondence between each retained residue and its position in the parent.

**Fig. 1.**
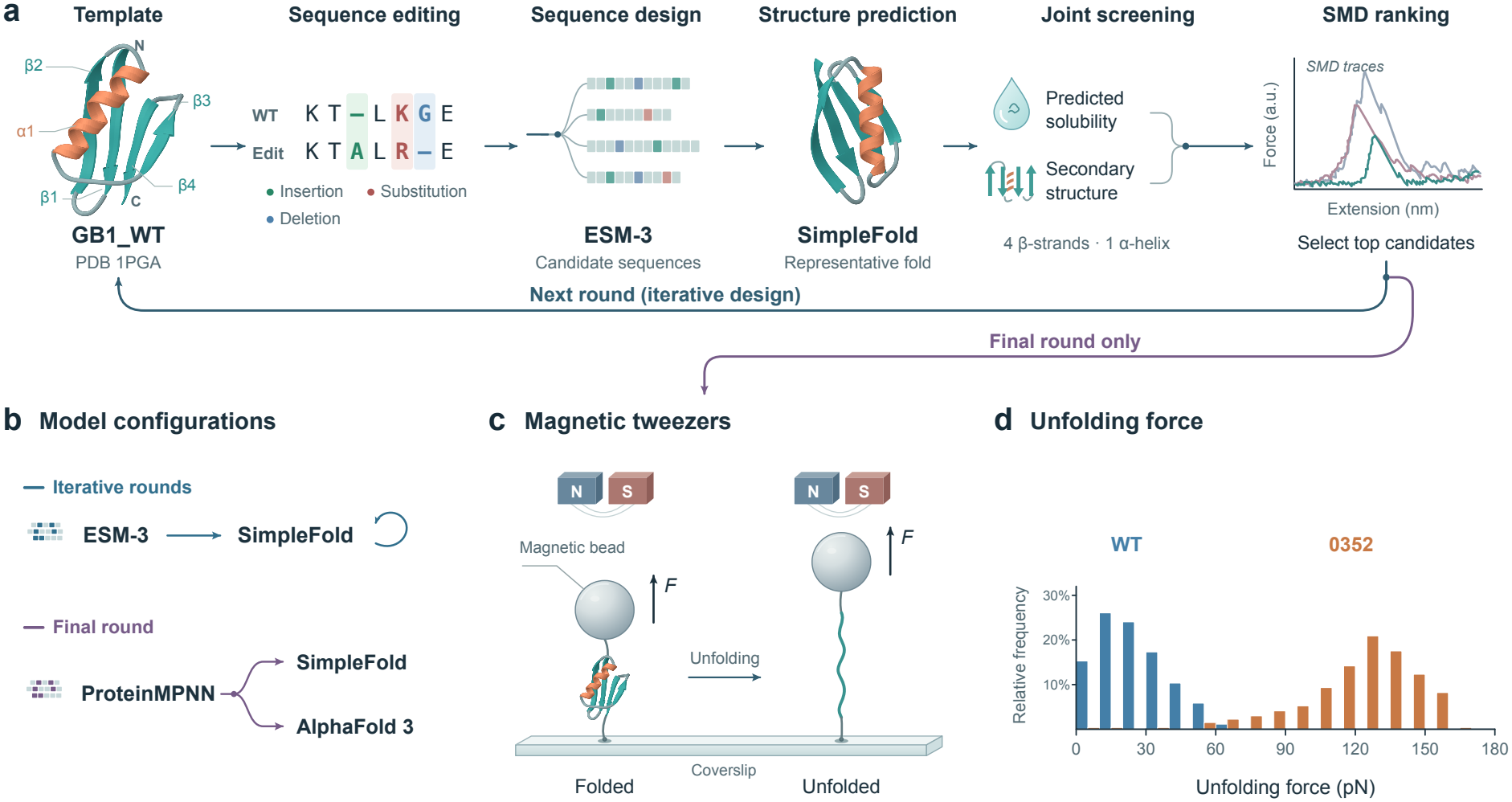
Iterative design and mechanical evaluation of GB1-like proteins. **a**, Sequence editing, fold prediction, joint screening and SMD ranking starting from wild-type GB1. The inset shows scaled force–extension traces. **b**, Models used during scaffold expansion and final-round sequence refinement. **c**, Magnetic-tweezers geometry. **d**, Example unfolding-force distributions for wild-type and design domains in the 98g3-0352 construct.

We compared five ensembles of about 2,930 unique sequences each (Fig. 1b and Supplementary Figs. 9– 12). One came from half-masked ESM-3 ^1^. For ProteinMPNN ^5^, we tested half and full masking at sampling temperatures *T* = 0.1 and 0.3. With half masking, median global identity to wild-type GB1 was *∼*60% for ESM-3 and *∼*80% for ProteinMPNN (Supplementary Figs. 9a and 10a). Across the 56 sites, ESM-3 had positional Shannon entropies of *∼*2.0–2.9 bits and wild-type recovery of *∼*0.5–0.7 (Supplementary Figs. 11 and 12). At some structurally constrained sites, ProteinMPNN recovered the wild-type residue almost every time, with entropy near zero. Elsewhere it sampled a wider range of residues. ESM-3 sequences had predicted isoelectric points (pI) near wild type and high Protein-Sol ^25^ scaled-sol values. Fully masked ProteinMPNN shifted the predicted pI towards *∼*10 and gave lower scaled-sol values, although SoluProt ^26^ predicted higher probabilities of soluble expression (Supplementary Fig. 9b–d). We used ESM-3 for changes in backbone length and ProteinMPNN for sequence refinement on a fixed backbone.

### Scaffold expansion and independent checks on the predicted fold

In four rounds of ESM-3 design, we expanded GB1 from 56 to 63, 75, 87 and 98 residues (rounds 63g1, 75g1, 87g1 and 98g1). At that length, ESM-3 redesigned selected sites in 98g2 and ProteinMPNN refined the sequence in 98g3 (Fig. 1a,b and Supplementary Table 1). Each batch shared coordinate masks and insertion anchors, with TM-score ^27^ and secondary structure guiding the search (Supplementary Table 1). We predicted structures locally with SimpleFold ^4^ for all 14,542 candidates. To pass the screen, a candidate had to meet the round-specific mean C*α* pLDDT threshold (*>*85 or *>*90), retain the required stretch of continuous DSSP ^28^ *β*-strand in each of four intervals and meet both the Protein-Sol ^25^ and SoluProt ^26^ thresholds (Supplementary Tables 2 and 3). The fast GROMACS screen at 50 nm ns^*−*1^ comprised 537 trajectories across 179 systems (Fig. 2c). In intermediate rounds, we pulled each shortlisted candidate once at 0.5 nm ns^*−*1^ and used the candidate with the highest peak force to seed the next round (Supplementary Table 3 and Supplementary Fig. 8).

**Fig. 2.**
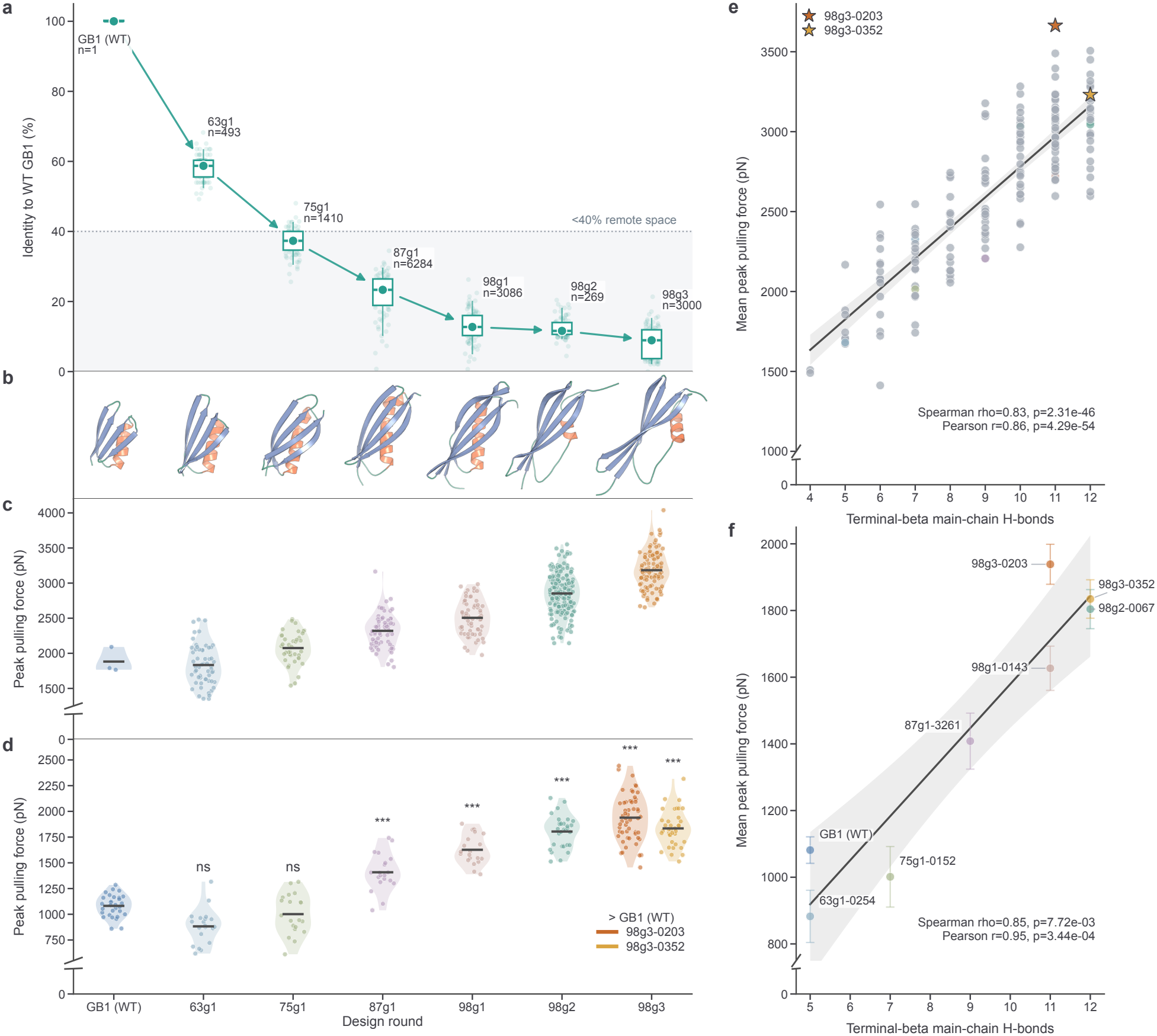
Scaffold expansion and simulated resistance to pulling. **a**, Global sequence identity to wild-type 1PGA. Circles, boxes and whiskers mark medians, interquartile ranges and 5th–95th percentiles of all sequences. Faint points are display subsamples. Arrows connect selected seeds, and the dotted line marks 40% identity. **b**, Representative structures from 56 to 98 residues. **c**, Raw trajectory maxima at 50 nm ns^*−*1^ (179 systems, 537 trajectories). **d**, Peak forces at 0.5 nm ns^*−*1^ after 20-point smoothing. From left to right, *n* = 30, 20, 20, 20, 20, 30, 55 and 37 trajectories. In c,d, points are trajectories, violins show distributions and ticks mark means. In d, one-sided Welch tests compare each variant with wild type, with Holm correction over seven comparisons (ns, adjusted *P ≥* 0.05 and ***, adjusted *P <* 0.001). **e**,**f**, Starting terminal-strand backbone hydrogen-bond count versus system mean peak force in c and d, respectively. Stars identify the two final designs. Error bars in f are 95% t-based confidence intervals on means. Lines are linear fits with 95% confidence bands. The panels report Pearson and Spearman correlations.

Global sequence identity to wild-type GB1 fell below the 40% reference line by rounds 75g1–87g1. Median identity reached 12.7% in 98g1 (*n* = 3,086), 11.6% in 98g2 (*n* = 269) and 8.9% in 98g3 (*n* = 3,000, Fig. 2a). The selected structures still had a predicted four-stranded *β*-sheet packed against an *α*-helix, now with a longer terminal clamp (Fig. 2b). SimpleFold assigned high mean confidence to the 87-residue ESM-3 seed 87g1-3261 and the 98-residue seeds 98g1-0143 and 98g2-0067. AlphaFold 3 ^3^ assigned lower confidence to parts of their expanded strands and core (Supplementary Fig. 2). In 98g3, ProteinMPNN generated 3,000 sequences on the fixed 98g2-0067 backbone at *T* = 0.3. Of the 182 that passed the combined screen, 19 also met the AlphaFold 3 criterion of mean C*α* pLDDT *>* 85 (10.4%, Supplementary Table 3). The two predictors agreed more closely for 98g3-0203 and 98g3-0352, the variants chosen for experiment (Supplementary Fig. 2).

At 0.5 nm ns^*−*1^, the first significant increase in mean peak force occurred in round 87g1 (Fig. 2d). We compared 20–55 trajectories per system against the wild-type NMR structure 3GB1 ^12^, which has the same sequence as 1PGA (Supplementary Table 5 and Supplementary Fig. 8). Wild-type GB1 reached 1081.4 *±* 106.5 pN (mean *±* SD). Neither early seed, 63g1-0254 or 75g1-0152, showed a significant increase over wild type in one-sided Welch t-tests with Holm correction (adjusted *P ≥* 0.05). From 87g1-3261 onward, all tested designs exceeded wild type (all adjusted *P <* 0.001, Fig. 2d). The highest final-round mean was 1938.7 *±* 221.9 pN for 98g3-0203 (*n* = 55), 1.79 times the wild-type mean. We selected a second high-ranking variant, 98g3-0352, that shared 50.0% sequence identity with 98g3-0203. It reached 1834.5 *±* 172.9 pN (*n* = 37), or 1.70 times the wild-type mean.

### Terminal-strand hydrogen bonds track force and cooperative clamp rupture

Pulling GB1 through its termini loads its parallel terminal *β*-strands in shear ^13–16^. We counted the main-chain hydrogen bonds between these strands in each starting structure and compared the count with that system’s mean peak force. Across 179 systems pulled at 50 nm ns^*−*1^, the correlations were Spearman *ρ* = 0.83 (*P* = 2.31 *×* 10^*−*46^) and Pearson *r* = 0.86 (*P* = 4.29 *×* 10^*−*54^, Fig. 2e). For the eight lineage representatives pulled at 0.5 nm ns^*−*1^, a 100-fold reduction in speed, Spearman *ρ* was 0.85 (*P* = 7.72 *×* 10^*−*3^) and Pearson r was 0.95 (*P* = 3.44 *×* 10^*−*4^, Fig. 2f). Wild-type GB1 and the early seeds had 5–6 bonds across the clamp. We counted 11 in 98g3-0203 and 12 in both 98g3-0352 and 98g2-0067. The mean peak force for 98g3-0203 lay above the fitted line, suggesting a contribution from contacts outside the terminal interface.

To identify which contacts broke near the force peak, we ran paired fixed-extension simulations starting 200 ps before and 200 ps after each peak (10 ns of NPT simulation at 310 K per rerun). We obtained 55 matched pairs for 98g3-0203 and 37 for 98g3-0352 (Fig. 3). Residue pairs with a mean occupancy loss of at least 0.40 clustered at the shear interface between the N-terminal strand (residues 1–24) and the C-terminal strand (residues 75–98, Fig. 3a). In one trajectory of 98g3-0352 (run 19), force rose while *∼*7–8 main-chain hydrogen bonds still spanned the core clamp (residues 9–18 and 82–93). The subsequent drop in force coincided with loss of contacts including Tyr7–Ile84 and Lys9–Asn86 (Fig. 3b). Across runs, the largest occupancy losses between residues 1–24 and 75–98 clustered along the terminal strand register (Fig. 3c). Summing over main-chain pairs assigned to *β*-strand (E–E) within residues 1–20 and 79–98 also gave lower occupancy after the peak in both designs (paired two-sided t-tests, Bonferroni-adjusted *P <* 0.001 for both, Fig. 3d).

**Fig. 3.**
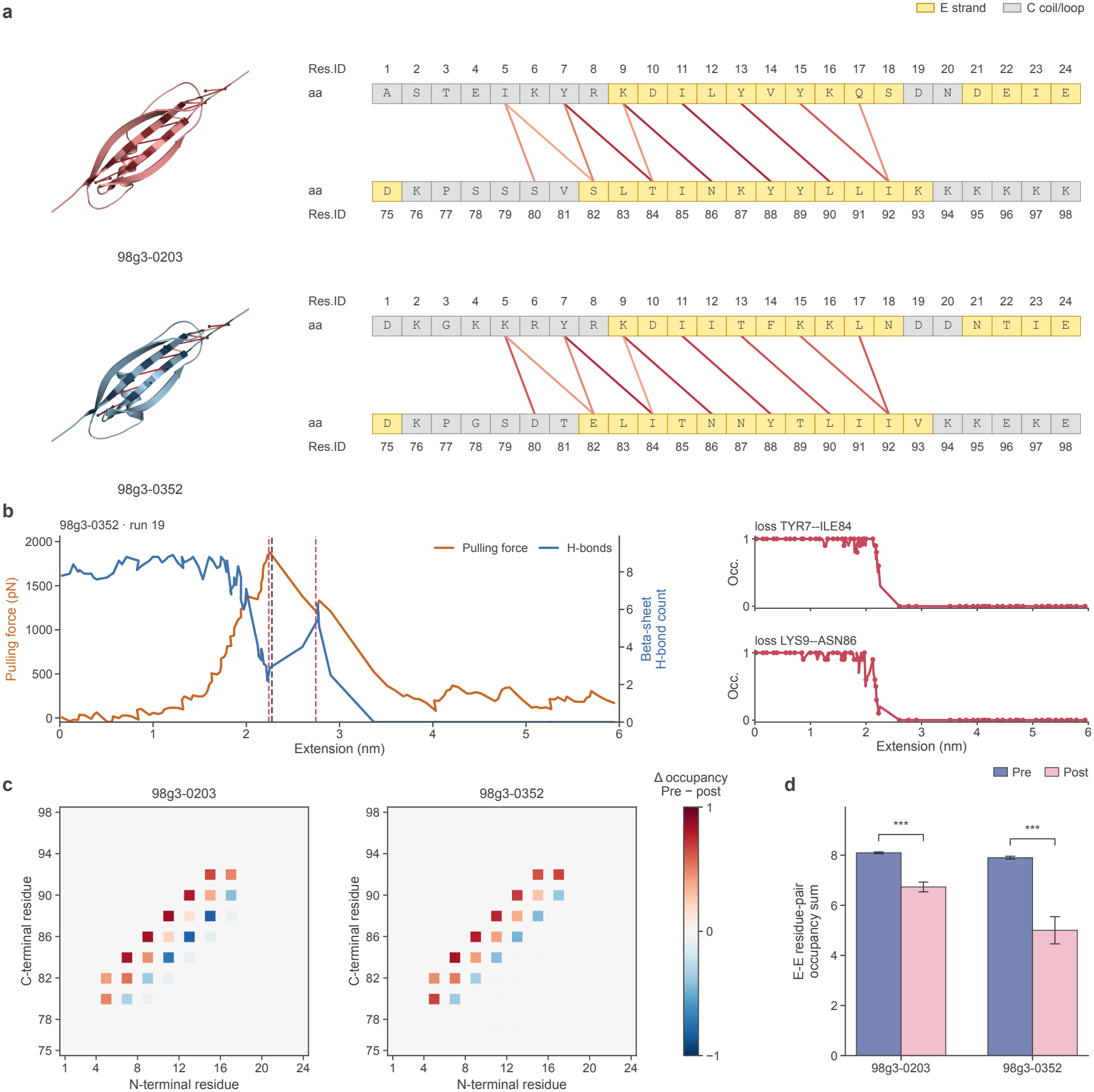
Terminal-strand hydrogen-bond occupancy around the force peak. **a**, Pairs with a mean occupancy loss of at least 0.4 are mapped onto representative post-peak structures. Strips show strand (E) and coil/loop (C) assignments in residues 1–24 and 75–98. **b**, Force and hydrogen-bond count (residues 9–18 versus 82–93) in pull 19 of 98g3-0352. Dashed lines locate the peak and rerun starting points. Insets show occupancy for two individual pairs. **c**, Mean pre-peak minus post-peak pair occupancy. Red indicates a decrease after rupture. **d**, Summed occupancies for selected main-chain pairs within the terminal *β*-strand regions. Bars show means *±* s.e.m. for 55 and 37 matched pre-/post-peak rerun pairs. Each fixed-separation rerun was initiated 200 ps before or after the peak. Paired two-sided t-tests with Bonferroni correction across the two designs gave adjusted *P <* 0.001 (***) for each.

### The fold at elevated temperatures and its predicted sensitivity to substitution

We ran one 500-ns unbiased MD trajectory for each protein at each temperature (310, 373, 430 and 450 K, Fig. 4 and Supplementary Figs. 5–7). From 310 to 430 K, all three proteins retained the four-strand, one-helix fold. The 56-residue wild type had a C*α* RMSD of *∼*0.1–0.2 nm. For the 98-residue designs, the corresponding ranges were *∼*0.2–0.4 nm at 310–373 K and *∼*0.3–0.6 nm at 430 K (Supplementary Figs. 5–7). The three trajectories diverged at 450 K. Wild-type 3GB1 lost much of its helix and sheet after *∼*400 ns, and its C*α* RMSD exceeded 2.5 nm (Fig. 4a,d,g). In 98g3-0203, the helix partially relaxed near 230 ns (*∼*1.2–1.3 nm RMSD), but the core sheet persisted and the final RMSD remained below that of unfolded 3GB1 (Fig. 4b,e,h). In 98g3-0352, the fold persisted throughout the 500 ns, with an RMSD of *∼*0.3–0.4 nm (Fig. 4c,f,i).

**Fig. 4.**
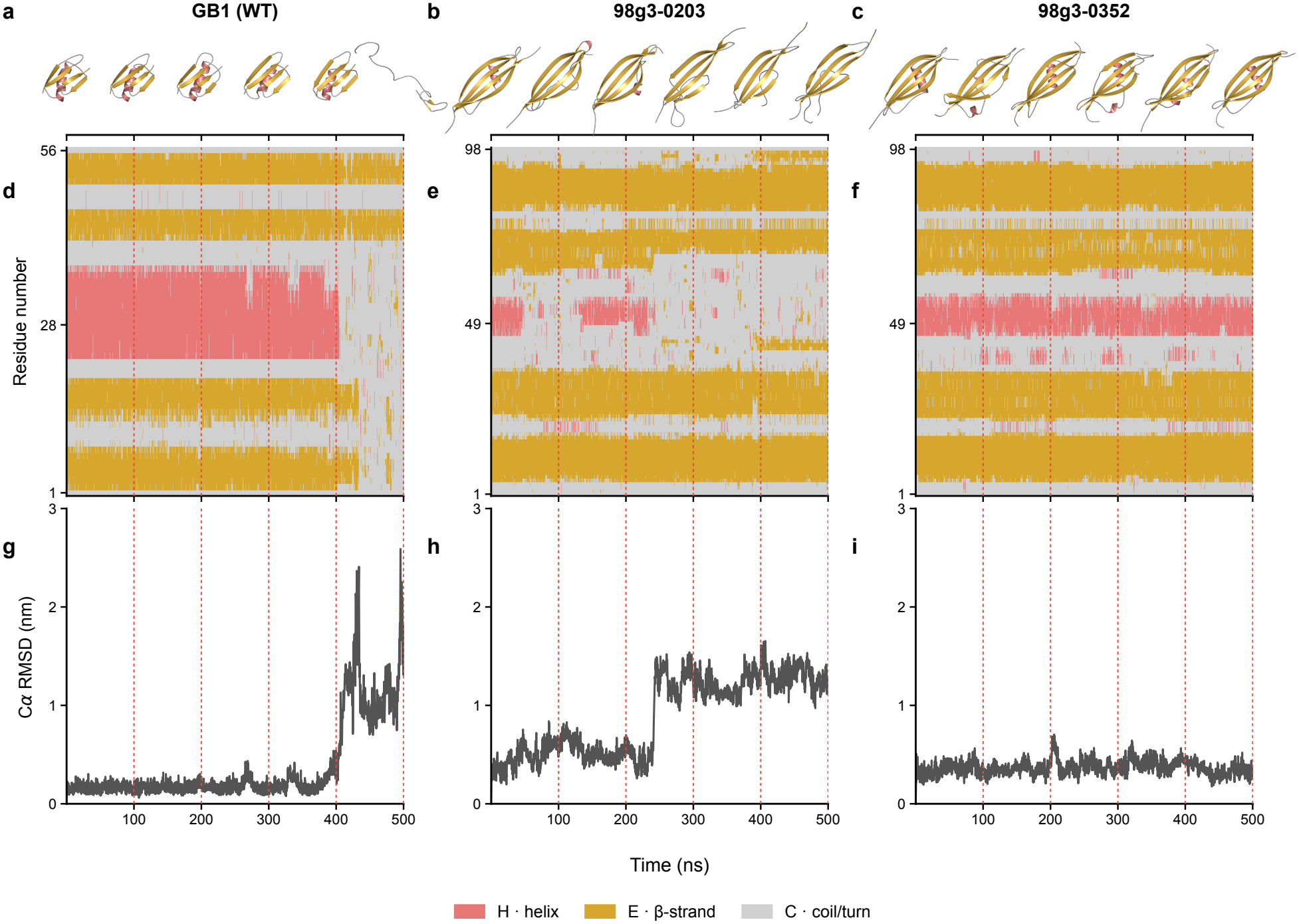
Structural retention in 500-ns trajectories at 450 K. **a–c**, Snapshots at 100-ns intervals for wild-type 3GB1, 98g3-0203 and 98g3-0352, respectively, fitted to the corresponding C*α* reference coordinates. **d–f**, Residue-by-time secondary structure for the same systems. DSSP assignments are grouped as helix (H), *β*-strand (E) and coil/turn (C). **g–i**, C*α* RMSD relative to the condition-specific starting structure after least-squares fitting. Red dashed lines indicate snapshot times. Each system contributed one trajectory sampled every 100 ps (5,001 frames).

We used ESM3ΔG ^29^, trained on large-scale cDNA-display proteolysis data ^30^, to estimate the effects of substitutions in GB1 and both designs (Supplementary Figs. 3 and 4). Across the expanded *β*-strands and core, the designs had higher predicted ensemble-mean per-residue ΔG values than wild-type GB1 (Supplementary Fig. 3a,b). The full scan covered all 20 amino acids at each site, giving 5,040 position-by-substitution values (Supplementary Fig. 4). At strand and hydrophobic core positions in both designs, the most destabilizing substitution had a large negative predicted ΔΔG (Supplementary Fig. 3d). Compared with wild type, the designs had few sites at which the maximum predicted ΔΔG exceeded +1 kcal mol^*−*1^ (Supplementary Fig. 3c).

### Contacts around the clamp and comparison with designed *β*-sheet proteins

We compared both final designs with wild-type 3GB1 and two 97-residue SuperMyo proteins, B42 and B347 ^21^, designed to maximize hydrogen bonding in the all-*β* immunoglobulin-like domain of titin I27 (Fig. 5 and Supplementary Table 4). The two GB1 designs shared 49 of 98 residues (50.0% identity). Their predicted structures both contained four *β*-strands and one *α*-helix and aligned closely (mTM-align ^31^ TM-score = 0.9297, RMSD = 1.149 Å). Relative to wild-type 3GB1, sequence identities were 9.9% for 98g3-0203 and 9.3% for 98g3-0352, with TM-scores of 0.6376 and 0.6409, respectively. Against B42 and B347, identities ranged from 18.9% to 25.2% and TM-scores from 0.4960 to 0.5187 (Fig. 5a and Supplementary Table 4).

**Fig. 5.**
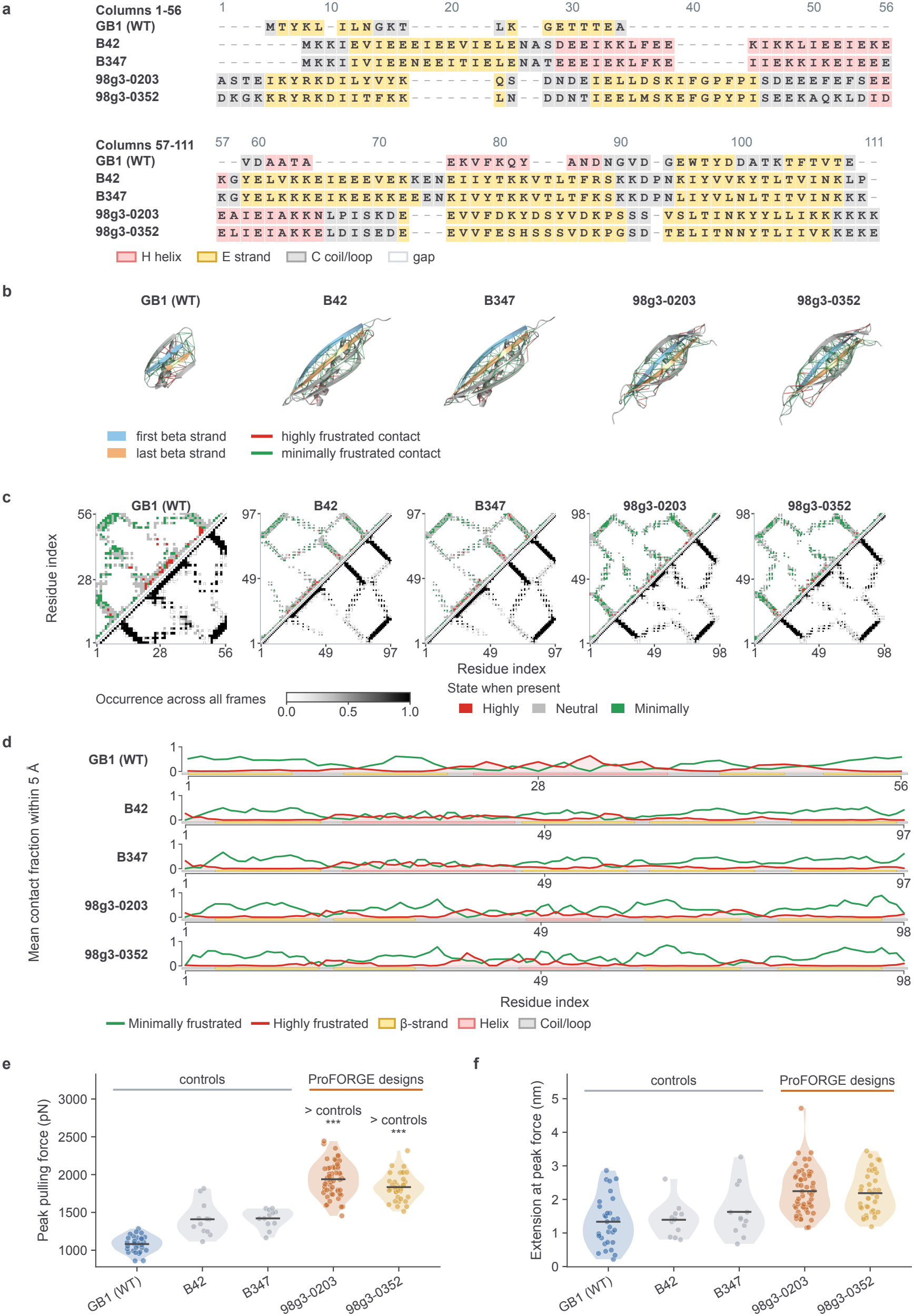
Comparison with wild-type GB1 and two designed *β*-sheet controls. **a**, PROMALS3D alignment coloured by DSSP secondary structure. **b**, Matched views of the terminal strands. Red lines mark highly frustrated contacts and green lines mark minimally frustrated contacts, each with all-frame state occupancy *≥*0.50. **c**, The upper triangle shows the most frequent frustration state while a contact was present. The lower triangle shows how often the contact occurred. **d**, Mean fractions of highly and minimally frustrated contacts within 5 Å of each C*α*. The denominator includes neutral contacts. **e**, Peak forces at 0.5 nm ns^*−*1^. From left to right, the five groups contain 30, 12, 12, 55 and 37 trajectories. One-sided Welch tests compare each design with each control, with Holm correction across all six comparisons. **f**, Extension at peak force. In e and f, each point represents one trajectory and horizontal ticks mark the means.

We calculated local configurational frustration ^32–34^ over equilibrium NPT trajectories for 3GB1, B42, B347, 98g3-0203 and 98g3-0352 (1,530, 612, 612, 2,805 and 1,887 frames, respectively, Fig. 5b–d). In both GB1 designs, minimally frustrated contacts with all-frame occupancy *≥* 0.50 linked the elongated terminal strands to the central helix and hydrophobic core (Fig. 5b,c). Within 5 Å of each C*α*, minimally frustrated contacts predominated along the load-bearing strands. Several loops and terminal regions had higher local fractions of frustrated contacts (Fig. 5d). In matched pulling simulations, both 98g3-0203 (*n* = 55) and 98g3-0352 (*n* = 37) had higher mean peak forces than 3GB1 (*n* = 30), B42 (*n* = 12) and B347 (*n* = 12). All six comparisons gave Holm-adjusted *P <* 0.001 in one-sided Welch t-tests (Fig. 5e). Extension at peak force averaged 2.25 *±* 0.68 nm for 98g3-0203 and 2.19 *±* 0.62 nm for 98g3-0352. Both means exceeded those of wild-type 3GB1 (1.33 *±* 0.74 nm), B42 and B347 (Fig. 5f).

### Magnetic tweezers resolve a fivefold increase in unfolding force

We measured the unfolding forces of 98g3-0203 and 98g3-0352 with single-molecule magnetic tweezers ^17,35^ (Fig. 1c,d and Fig. 6a). Each polyprotein contained two copies of one design flanked by four wild-type domains (DogTag–GB1–GB1–design–design–GB1–GB1–SpyTag). Covalent attachment to a DogCatcher-coated magnetic bead ^36^ and a SpyCatcher-coated coverslip ^37^ put all six domains under the same N-to-C-terminal load and force calibration (Fig. 6a,b). We recorded 178 cycles across five recordings for 98g3-0203 and 148 across three for 98g3-0352, loading at a nominal 1 pN s^*−*1^ and unloading at 0.2 pN s^*−*1^. The traces showed small steps from wild-type GB1 at *∼*15–35 pN followed by larger steps from the 98-residue domains at forces above 90 pN (Fig. 6c).

**Fig. 6.**
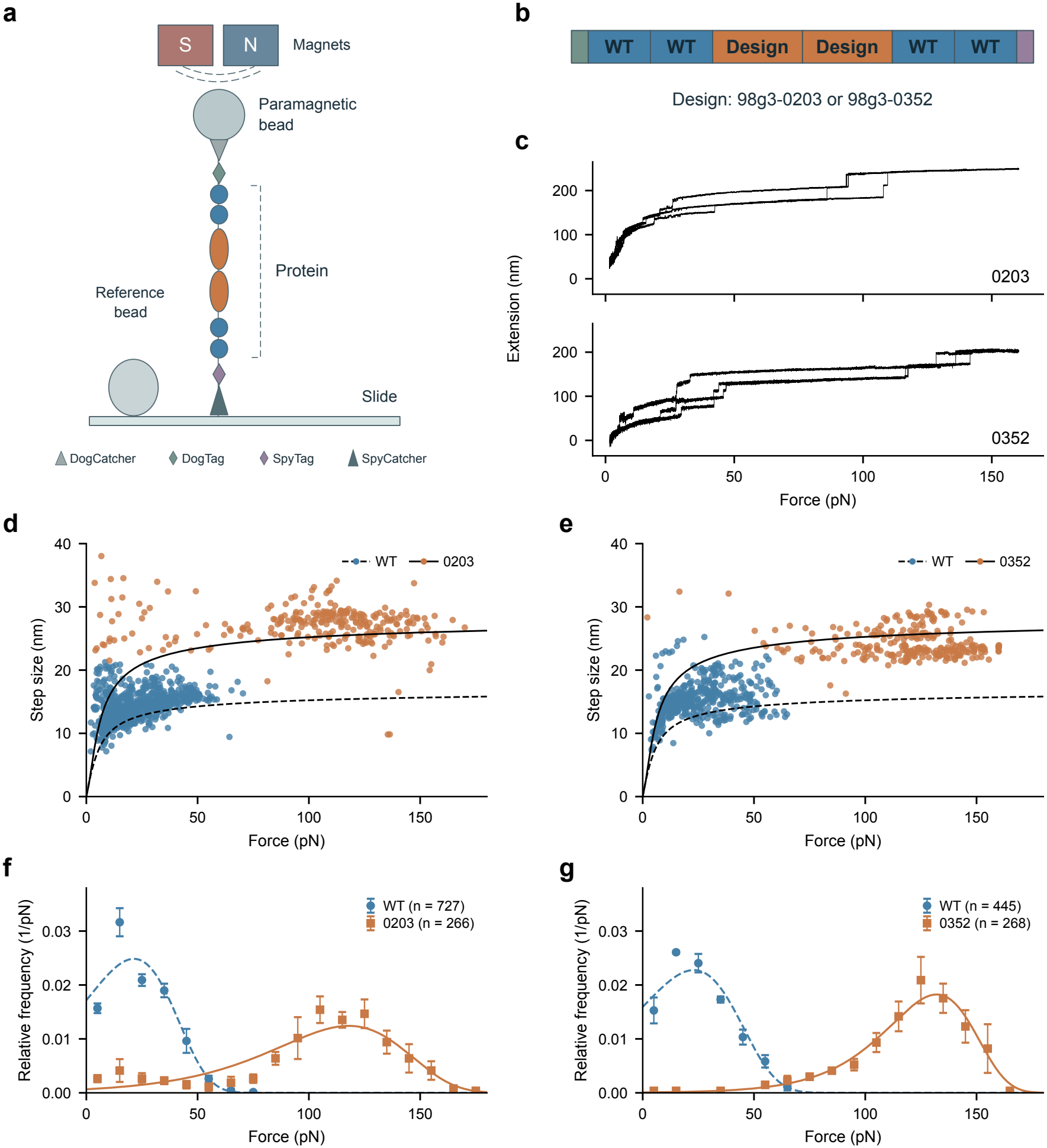
Single-molecule unfolding forces in mixed-domain tethers. **a**,**b**, Magnetic-tweezers geometry and domain order. Four wild-type GB1 domains flank two copies of the design. **c**, Cycles 8–10 from one recording per construct, at a nominal loading rate of 1 pN s^*−*1^. **d**,**e**, Step size versus force for the 98g3-0203 and 98g3-0352 constructs. Blue denotes wild-type events and orange denotes design events. The dashed and solid curves show the respective polymer-model predictions. **f**,**g**, Unfolding-force densities in 10-pN bins. Each point gives the fraction of events in a bin divided by its width. Error bars show one bootstrap standard error from resampling whole recordings. Curves show the fitted Bell distributions. Five recordings of 98g3-0203 yielded 727 wild-type and 266 design events. Three recordings of 98g3-0352 yielded 445 wild-type and 268 design events.

We assigned events by force and step size using a bivariate Gaussian mixture. The measured steps followed the worm-like chain / freely jointed chain (WLC–FJC) curves ^38,39^ for 56-residue wild-type GB1 (folded length *b* = 2.65 nm) and the 98-residue designs (*b* = 6.06 nm for 98g3-0203 and 6.00 nm for 98g3-0352, Fig. 6d,e). We fitted the constant-loading-rate Bell distribution ^17^ by maximum likelihood to estimate the most probable unfolding force (*F* ^*\**^). For 98g3-0203, this force was 117.9 pN (266 events), or 5.5 times the 21.4 pN measured for its internal wild-type control (727 events, Fig. 6f). For 98g3-0352, it was 132.1 pN (268 events), or 5.8 times the wild-type value of 22.9 pN (445 events, Fig. 6g). The wild-type-to-design event ratios were 2.7:1 and 1.7:1, respectively, for tethers with four wild-type and two design domains.

The designs ranked in opposite orders in simulation and experiment. At 0.5 nm ns^*−*1^, 98g3-0203 had the higher mean peak force in SMD (1938.7 versus 1834.5 pN, Fig. 2d). At 1 pN s^*−*1^ in the magnetic tweezers, 98g3-0352 had the higher most probable unfolding force (132.1 versus 117.9 pN, Fig. 6f,g). We also observed a small group of large steps below 50 pN for 98g3-0203 (Fig. 6d,f). Such events were rare for 98g3-0352 (Fig. 6e,g) and could reflect conformational heterogeneity or incomplete refolding. In the starting structure, 98g3-0203 had a C-terminal polylysine stretch (residues 93–98, Fig. 3a and Supplementary Table 6) and 11 terminal hydrogen bonds, compared with 12 in 98g3-0352 (Fig. 2f). In the 500-ns trajectories at 450 K, the helix partially relaxed in 98g3-0203 and the fold persisted in 98g3-0352 (Fig. 4c,e,f,h,i). Measurements at several loading rates could establish whether the ordering depends on rate. Varying the refolding interval between ramps would help establish whether incomplete refolding accounts for the low-force events.

## Discussion

In the GB1 lineage, following the terminal strands through each insertion made it possible to compare the same mechanical feature across scaffolds of different lengths (Fig. 1 and Supplementary Fig. 1). The number of backbone hydrogen bonds spanning these strands tracked the mean peak force at two simulated pulling speeds a factor of 100 apart (Fig. 2e,f). The bonds counted before pulling lie at the interface where contacts were lost near the force peak (Fig. 3). Their number is therefore a useful descriptor for this loading geometry, although the rest of the fold can also contribute to the force needed for rupture.

Half-masked ESM-3 sampled divergent sequences, while ProteinMPNN repeatedly selected the same amino acids at some constrained sites (Supplementary Figs. 9–12). That difference guided our use of ESM-3 to expand the scaffold and ProteinMPNN to refine it once the backbone was fixed. Independent structure prediction narrowed the final-round set from 182 locally screened sequences to 19 that met the AlphaFold 3 confidence cutoff. The other 163 were excluded on predicted confidence alone. Their folding behaviour remains unknown. We measured two of the selected designs by magnetic tweezers.

Extending GB1 from 56 to 98 residues roughly doubled the number of hydrogen bonds across the terminal *β*-clamp (Fig. 2e,f). Contacts at this interface broke near the simulated force peak (Fig. 3), and minimally frustrated contacts linked the strands to the core (Fig. 5b–d). B42 and B347, two SuperMyo designs derived from titin I27, extend an all-*β* fold through backbone diffusion followed by ProteinMPNN sequence design ^21^. For GB1, residue mapping and strand-continuity filters kept track of the elongated interface within a predicted mixed *α*/*β* fold. Under matched simulated pulling, both final GB1 designs had higher mean peak forces than B42 and B347 (Fig. 5e).

The GB1 experiments establish what this design strategy achieved. Despite sharing less than 10% global sequence identity with wild-type GB1, both final designs retained the terminal clamp in their predicted structures. At a nominal loading rate of 1 pN s^*−*1^, the most probable unfolding force was 117.9 pN for 98g3-0203 and 21.4 pN for its wild-type control. For 98g3-0352, the corresponding values were 132.1 and 22.9 pN (Fig. 6f,g). Within their respective tethers these gains were 5.5-fold and 5.8-fold. Both designs exceeded wild type in simulation and experiment, but their relative order reversed. Iterative selection on simulated force thus identified mechanically resistant domains. Their ordering at a specified loading rate still requires experimental calibration.

These two force ranges suggest a specific use for the designs as building blocks in protein materials. GB1 polyproteins can act as molecular springs ^14^, and experiments with GB1-based hydrogel crosslinkers show that the surrounding polymer network influences whether the domains unfold under strain ^40^. Pairing wild-type GB1 with a designed domain in a tandem crosslinker could allow extension in two stages as the load rises. A direct test would compare gels containing mixed crosslinkers with gels containing only wild-type or only designed domains at the same crosslink density. Cyclic stress–strain measurements would establish whether the separation seen in a single tether changes the energy dissipated during stretching or the recovery after unloading.

The same separation could also be useful in a single-molecule force probe. Wild-type GB1 and the longer designs produce steps that differ in size as well as force, offering two ways to identify which domain has unfolded. Calibrating those distributions over a range of loading rates and refolding intervals would establish where the two signals remain distinguishable in repeated use. A separate test of the computational method would be to follow a load-bearing interface in another fold or to change the pulling geometry within GB1. Comparing the resulting bond counts and measured unfolding forces would test whether the relationship found for terminal shear in GB1 carries over to a different load path.

## Methods

### Workflow construction and sequence generation

We ran ProFORGE locally, importing the source PDB chain together with its sequence and residue indices. Protein Studio recorded the sequence and coordinate masks, insertion and deletion sites, secondary structure and residue-conservation constraints in the prompt. After each edit, the software mapped retained positions back to the parent sequence. We started design from the GB1 crystal structure 1PGA ^11^ and used the NMR structure 3GB1 as the wild-type control for pulling simulations ^12^.

For local ESM-3 inference we used esm3-sm-open-v1. Within each batch in the first four rounds, all sequences shared one positional editing plan. The search assigned unit weight to the TM-score relative to the round’s seed ^27^ and to agreement with the prompted secondary structure. All other weights were zero. For round 98g2, we chose 17 positions independently in each sequence for redesign and retained the other 81. In 98g3, ProteinMPNN redesigned the full sequence on the SimpleFold-predicted 98g2-0067 backbone at a sampling temperature of 0.3. Supplementary Table 1 lists the parents, permitted edits and sequence counts for each round. Supplementary Table 6 gives the seven reported design sequences.

For the generator comparison, half-masked ESM-3 and half-masked ProteinMPNN at temperatures 0.1 and 0.3 each yielded 2,930 unique sequences. Fully masked ProteinMPNN yielded 2,927 at 0.1 and 2,930 at 0.3. We counted each unique sequence once. At each site, we calculated wild-type recovery as the fraction of sequences carrying the wild-type residue and entropy in bits from the observed amino-acid frequencies. For the half-masked panels in Supplementary Fig. 12, we subtracted 0.5 from the wild-type frequency, set negative values to zero and renormalized the frequencies at that site. The fully masked panels show the observed frequencies.

### Structural screening and sequence comparison

For each sequence, SimpleFold 3B generated one structure in 500 steps at *τ* = 0.05. We first screened on the mean C*α* predicted local distance difference test (pLDDT) score. In each of four prespecified intervals, we used DSSP ^28^ to find the longest uninterrupted run of *β*-strand (E) and required it to meet the minimum length for that generation. The remaining sequences had to pass thresholds for both Protein-Sol scaled-sol ^25^ and the SoluProt v1.0.1 probability of soluble expression ^26^. Supplementary Tables 2 and 3 give the cutoffs and counts. We submitted all 182 final-round sequences that passed these filters to AlphaFold 3 and retained those with mean C*α* pLDDT *>* 85.

We calculated global sequence identity with EMBOSS Needle using the Needleman–Wunsch algorithm ^41^, the EBLOSUM62 matrix and gap-opening and extension penalties of 10.0 and 0.5. For the lineage comparisons we used 1PGA as the reference and disabled end weighting. All identity percentages use the full alignment length, including gaps. The seven pairwise comparisons centred on the final designs included 3GB1, B42 and B347, with end-gap parameters of 10.0 and 0.5. We obtained the pairwise TM-scores and RMSDs from one mTM-align v20180725 alignment^31^ (Supplementary Table 4).

### Steered molecular dynamics and seed selection

We prepared systems for GROMACS pulling simulations ^23,24^ with CHARMM36m ^42^, TIP3P water ^43^, neutralizing ions and 0.15 M NaCl. We restrained the N-terminal C*α* atom and pulled the C-terminal C*α* atom along +z over a nominal 10 nm. The pulling spring constant was 1,000 kJ mol^*−*1^ nm^*−*2^. The fast screen used 50 nm ns^*−*1^ and the detailed and production simulations used 0.5 nm ns^*−*1^. Supplementary Table 5 gives equilibration, pulling and output-sampling parameters.

We ranked fast pulls by the maximum raw force magnitude in each trajectory. The fast screen covered 179 systems including wild type, with 537 trajectories in total. In 98g1, 24 sequences passed the combined screen and 21 systems entered fast pulling. In 98g3, the 19 AlphaFold 3 passes corresponded to 18 unique sequence identifiers and 36 simulation systems. We ran one detailed pull for each shortlisted high-force candidate and used the candidate with the highest raw maximum to seed the next round. For replicate comparisons, we pooled the detailed trajectories with additional runs at the same physical parameters. We averaged the force magnitude over a moving window of 20 points and took the maximum of the smoothed trace for each trajectory.

### Terminal-strand hydrogen bonds

For each system we took the starting coordinates from one representative run and identified the first and last continuous *β*-strands with PyMOL dss. We counted backbone nitrogen-donor/oxygen-acceptor pairs between these strands using gmx hbond in GROMACS 2025.3. A hydrogen bond required a donor–acceptor distance *≤*0.35 nm and an acceptor–donor–hydrogen angle *≤*30^*°*^. The raw output distinguished individual hydrogens. For the static bond count, we merged events with the same donor and acceptor.

For the two final designs, we initialized fixed-separation NPT reruns 200 ps before and 200 ps after each production force peak. A harmonic restraint centred at the corresponding terminal separation along the pulling axis was held fixed for 10 ns at 310 K (spring constant 1,000 kJ mol^*−*1^ nm^*−*2^). We analysed 10,001 frames per rerun at 1-ps spacing. Residue-pair occupancy was the fraction of frames containing at least one inter-residue main-chain hydrogen bond. For Fig. 3d we summed occupancies over pairs whose two residues lay within the union of residues 1–20 and 79–98 and were assigned *β*-strand in the reference annotation. The set included pairs within a terminal region and pairs linking the two regions.

### Thermal simulations and stability predictions

We simulated 3GB1 and each final design for 500 ns at 310, 373, 430 and 450 K, with one trajectory per protein per temperature. We used GROMACS 2025.3, the CHARMM36-jul2022 force field, TIP3P water, 0.15 M NaCl and neutralizing ions in a 5.0 *×* 5.5 *×* 9.0 nm box. Simulations used a 2-fs step, velocity-rescaling temperature control and an isotropic Parrinello–Rahman barostat at 1 bar. Coordinates were saved every 100 ps. C*α* RMSDs were calculated after least-squares fitting to each starting structure. DSSP assignments were grouped into helix, strand and coil/turn states.

ESM3ΔG predictions ^29,30^ were averaged across three model repeats per protein. Predicted substitution effects were calculated as ΔΔ*G* = *G*_mutant_ *− G*_reference_, using the input sequence as the reference at each site. The full matrix included all 20 amino acids at each of 252 positions across the three proteins. The extrema in Supplementary Fig. 3 use the 19 non-reference amino acids per position.

### Configurational frustration in equilibration ensembles

For each NPT equilibration trajectory, we sampled 51 frames at 10-ps intervals from 500 to 1,000 ps. This gave 1,530 frames for 3GB1, 612 each for B42 and B347, 2,805 for 98g3-0203 and 1,887 for 98g3-0352. FrustratometeR ^32–34^ classified contacts as highly frustrated (index *≤ −*1), neutral (*−*1 *<* index *<* 0.78) or minimally frustrated (index *≥* 0.78). To calculate contact occurrence, we divided the number of frames containing a pair by the total number of frames. For occupancy in a given state, we counted only frames where the pair occupied that state and used the same denominator. Conditional state fractions instead used the number of frames containing the pair as their denominator. For each residue in each frame, we calculated the fraction of contact midpoints within 5 Å of its C*α* that belonged to each state. We averaged those fractions over frames, assigning zero when no contact lay within the radius.

### Magnetic-tweezers measurements and event assignment

Each tether contained DogTag–GB1–GB1–design–design–GB1–GB1–SpyTag. We coupled the DogTag end to a DogCatcher-coated paramagnetic bead ^36^ and the SpyTag end to a SpyCatcher-coated coverslip ^37^. Both constructs were pulled through their N- and C-termini. *[Expression, purification and sample-preparation details for the two constructs are pending author confirmation*.]

We followed the magnetic-tweezers setup and calibration of Guo et al. ^17^. The instrument used an Olympus IX73 microscope with a 100*×* oil-immersion objective, a PIFOC Nano focusing actuator, an LS-110 motorized stage, two permanent magnetic rods and a XIMEA MQ022MG-CM camera. We estimated extension from bead images using a focal-plane library sampled at 50-nm intervals and corrected drift with an immobilized reference bead. The reported extension resolution was *<*2 nm. Force calibration combined Stokes drag measurements on M270 beads in 90% glycerol with a bead-specific transverse-fluctuation measurement near 10 pN. Estimated force accuracy was about 5%. Recordings with temperature annotations specified 24.2–24.4 ^*°*^C. For magnet distance d in millimetres, the force in piconewtons was *C*[exp(*−d/*0.81799) + 1.82384 exp(*−d/*0.29115)], with a separate C for each recording. For the 6 November B1 recording of 98g3-0203, we obtained *C* = 119.99 from the median ratio of the instrument-reported force to the calibration function evaluated at *C* = 1.

We used nominal loading and unloading rates of 1 and 0.2 pN s^*−*1^, respectively, and analysed up to the first 50 loading cycles in each recording. We detected candidate steps on 0.1-s median extension traces, then fitted them to the original samples with a local slope. For fitted amplitudes *≥*7 nm, we located the step using the median of ten samples on either side. After trace review, we retained steps of 7–40 nm and calculated the force from the magnet distance at the original sample marking the upward step.

Within each construct we standardized force and step size separately, then fitted a two-component bivariate Gaussian mixture with a full covariance matrix for each component. Expectation–maximization used 64 initializations, up to 1,000 iterations and a convergence tolerance of 10^*−*7^. Each event was assigned to the component with higher posterior probability. The components with lower and higher mean forces were assigned to wild-type and design domains, respectively. The theoretical step was the unfolded worm-like-chain extension ^38^ minus the mean projection of the folded domain, treated as a freely rotating rigid rod ^39^. We used 0.36 nm contour length per residue, a persistence length of 0.8 nm and *k*_B_*T* = 4.1 pN nm. Folded lengths were 2.65, 6.06 and 6.00 nm for wild type, 98g3-0203 and 98g3-0352, respectively.

### Statistical analysis

For SMD, we report the mean and standard deviation of the peak forces across trajectories. We used one-sided Welch t-tests to compare each design with its comparator and applied Holm correction separately to the seven design–wild-type comparisons in Fig. 2d and the six design–control comparisons in Fig. 5e. For correlations between hydrogen-bond count and force, each system contributed one observation to the Pearson and Spearman coefficients. In the fixed-separation reruns, we compared summed occupancies before and after the peak using a paired two-sided t-test within each design, with Bonferroni correction across the two designs. We report descriptive statistics for extensions at peak force, the generator ensembles and the trajectories at four temperatures.

For each construct, we fitted the constant-loading-rate Bell distribution ^17^ by maximum likelihood to the individual event forces for each domain type, using the median measured loading rate for each recording. We weighted each recording’s density by its event count. The force at the maximum of the combined density gave the most probable unfolding force. To estimate histogram error bars, we drew 2,000 bootstrap samples per construct, resampling whole recordings with replacement so that events from a given recording stayed together.

## Data availability

The seven reported sequences are given in Supplementary Table 6. Screening counts, structural comparisons and simulation parameters are given in Supplementary Tables 1–5. *[Add the data repository and accession or DOI for the simulation outputs, magnetic-tweezers traces, recording-specific calibrations, assigned events and histogram counts*.]

## Code availability

The source code for ProFORGE, the platform used for protein design and structural screening in this study, is publicly available at https://github.com/natsusorahoshinochan-max/modules-ESM. The repository includes installation instructions, documentation for Protein Studio and the workflow modules, and example modules. Model inference requires local model weights or credentials for the selected external service, depending on the configuration.

## Acknowledgements

This work was supported by the National Natural Science Foundation of China (U24A2014) and the Wenzhou Science and Technology Bureau (ZY2024002). Wanqing Yang received support from the Graduate Scientific Research Foundation of Wenzhou University (3162025003042). Yang Wang received support from the Natural Science Foundation of Zhejiang Province (MS27A040001) and the Natural Science Foundation of Wenzhou (2025G0009). Yanwei Wang received support from the National Natural Science Foundation of China (12674266).

## Author contributions

Yanzhang Huang, Wanqing Yang and Yang Wang contributed equally to this work. Yanzhang Huang and Yang Wang led the computational work, designing GB1 variants and using molecular dynamics simulations to characterize their response to force. Wanqing Yang led the single-molecule magnetic tweezers experiments to measure the unfolding forces of the designed proteins, with contributions from Zilong Guo and Jie-Li Wang. Jianwei Shuai and Fangfu Ye jointly supervised the protein design and simulations, and Yanwei Wang and Hu Chen jointly supervised the single-molecule experiments.

