## Supplemental Data 1 for "ProFORGE: a composable sequence–structure–dynamics workflow validated by mechanoprotein design"

### Supplementary figures

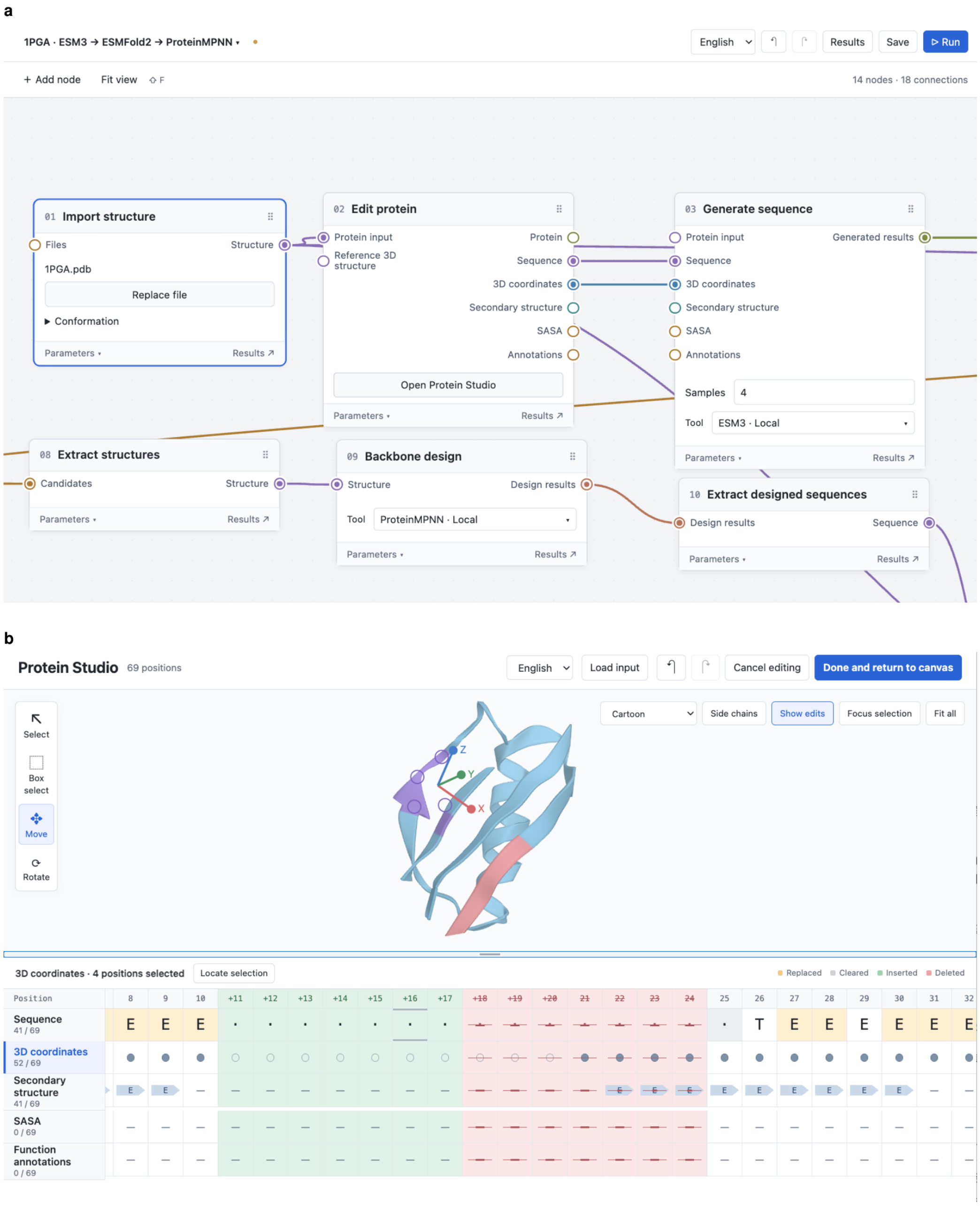


**Supplementary Fig. 1: ProFORGE software interface.** (a) Partial view of the node-based workflow editor, showing node connections, tool selection, and access to parameters. (b) Protein Studio, showing the protein structure, per-position information tracks, and editing states.

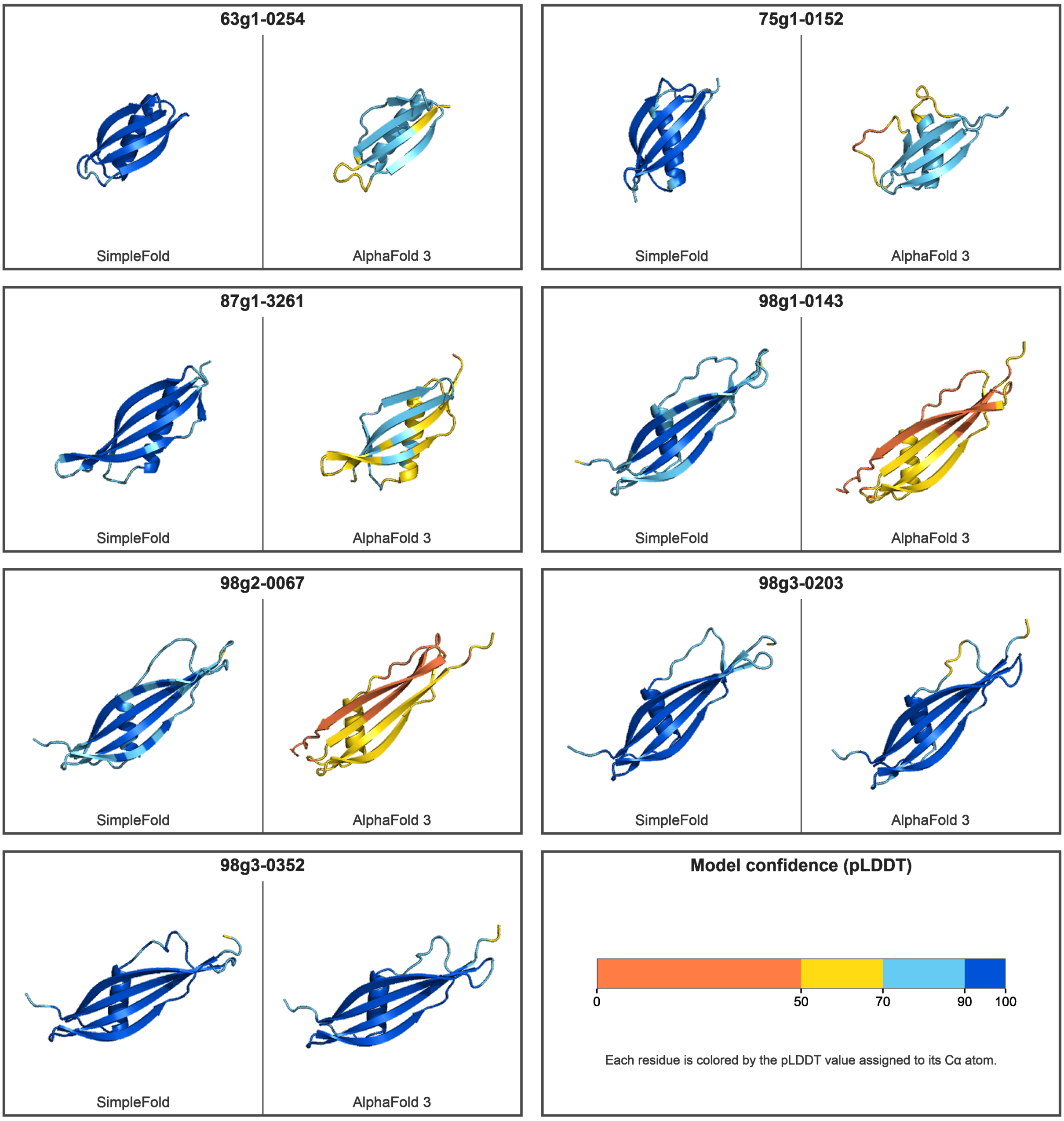


**Supplementary Fig. 2: Paired SimpleFold and AlphaFold 3 predicted structures for seven lineage designs.** For each design, the SimpleFold prediction is on the left and the AlphaFold 3 prediction is on the right. From top to bottom, the rows contain 63g1-0254 and 75g1-0152; 87g1-3261 and 98g1-0143; 98g2-0067 and 98g3-0203; and 98g3-0352, followed by the pLDDT color key. Each residue is colored by the pLDDT value assigned to its C$\alpha$ atom. Structures are shown at a common scale.

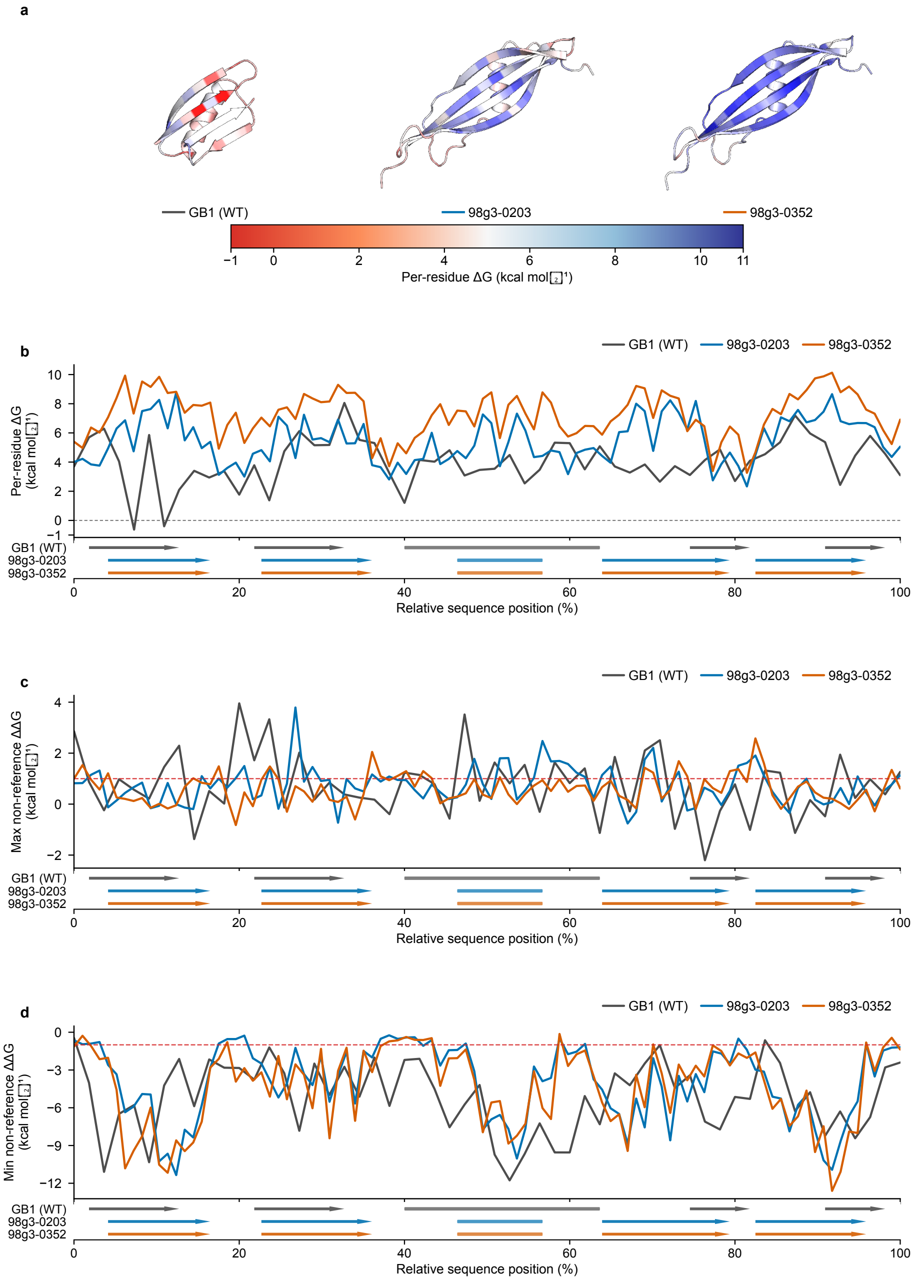


**Supplementary Fig. 3: Per-residue ESM3ΔG and non-reference amino-acid substitution summaries.** (a) Structures colored by ensemble mean per-residue ΔG, with lower values in red and higher values in blue. (b) Ensemble mean per-residue ΔG against relative sequence position. (c) Maximum ΔΔG across the 19 non-reference substitutions at each position, with a dashed guide at $+1$ kcal mol$^{-1}$. (d) Minimum non-reference ΔΔG, with a dashed guide at $-1$ kcal mol$^{-1}$. Secondary-structure strips use arrows for beta strands and rounded bars for alpha helices. Gray, blue, and orange denote GB1 (WT), 98g3-0203, and 98g3-0352, respectively. Predictions are ensemble means of three model repeats per protein.

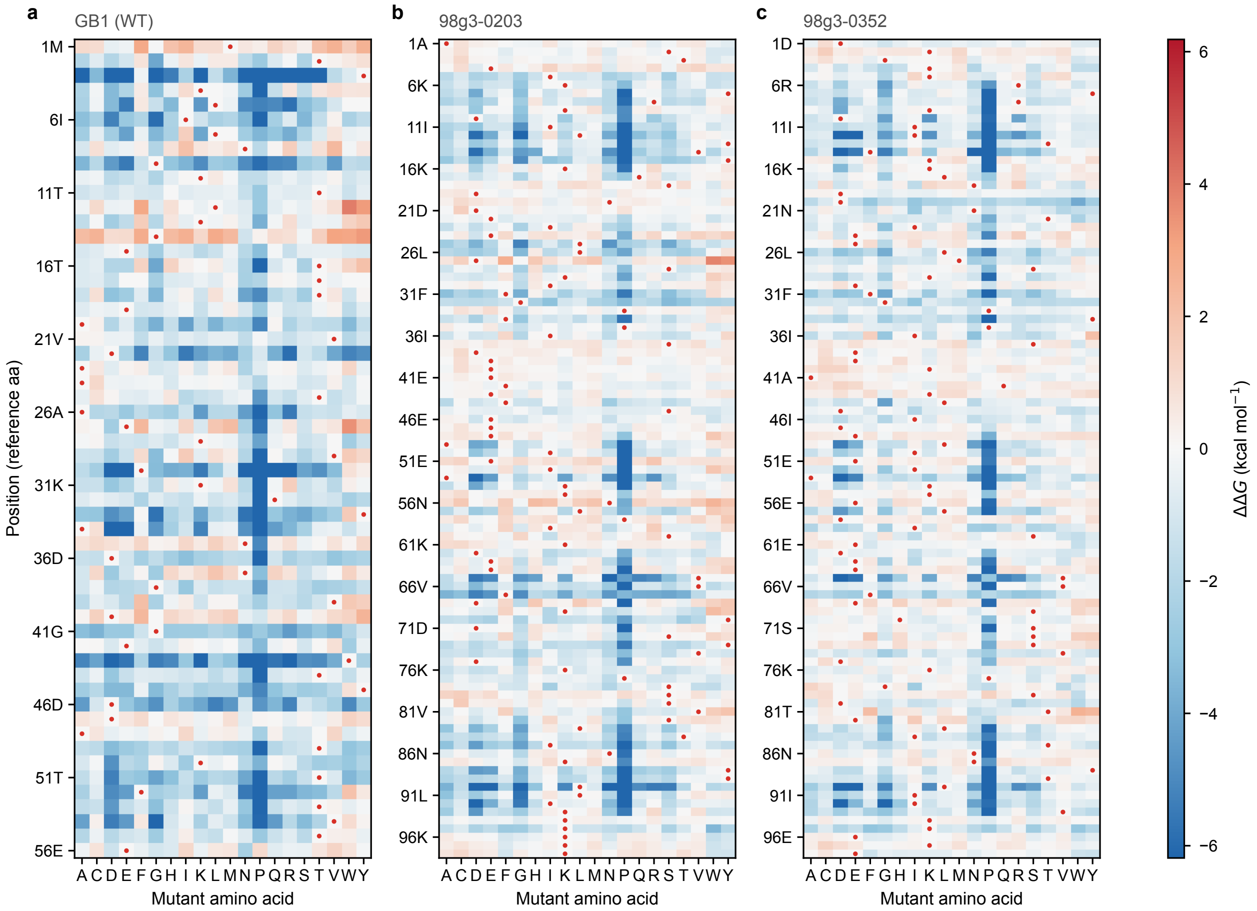


**Supplementary Fig. 4: Signed ESM3ΔG amino-acid substitution matrices.** Heatmaps show ensemble mean ΔΔG for substitution with each of the 20 standard amino acids at every residue position in (a) GB1 (WT), (b) 98g3-0203, and (c) 98g3-0352. Red dots mark the reference amino acid. Negative values are blue and positive values are red. All panels share a symmetric color scale whose limits are defined by the 98th percentile of |ΔΔG| across the complete three-system matrix. The complete position-by-substitution matrix contains 5,040 cells.

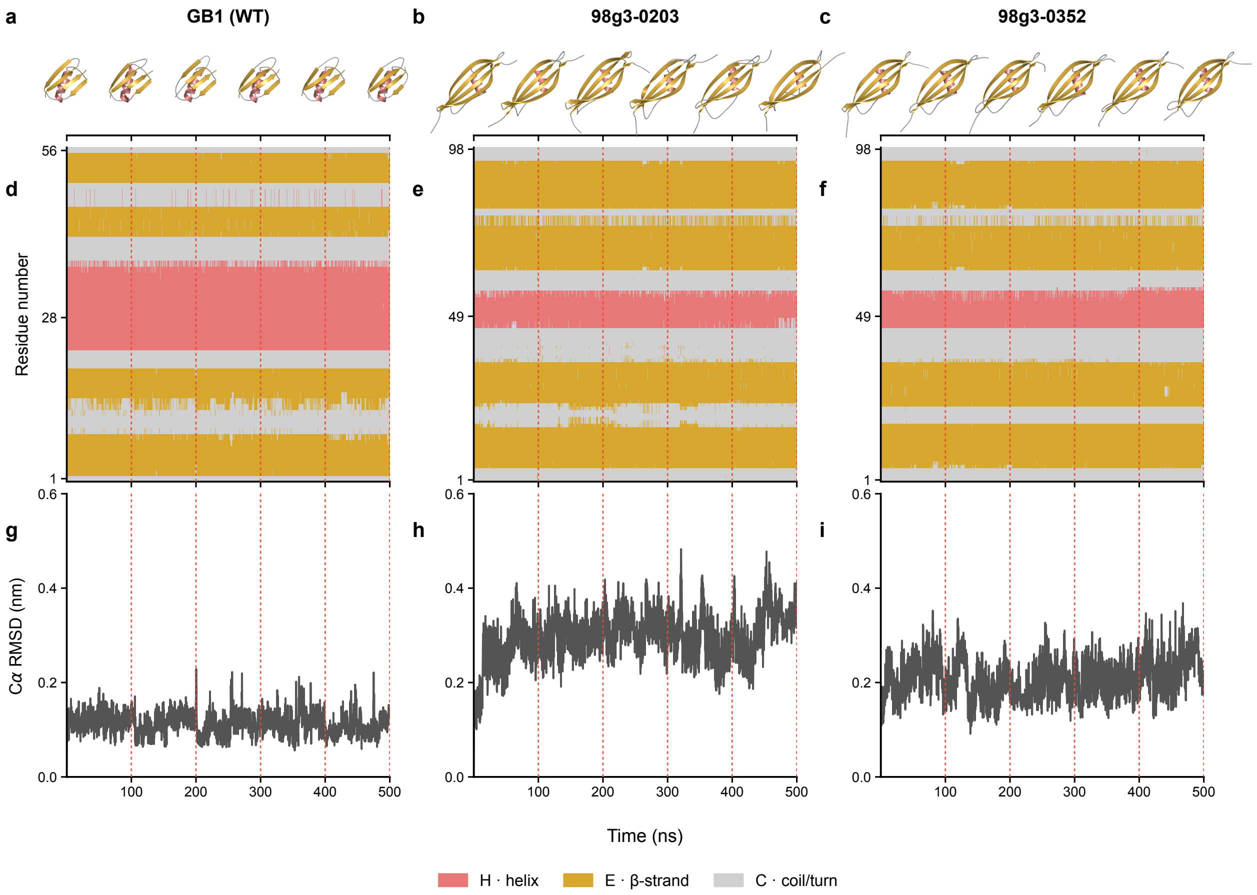


**Supplementary Fig. 5: Structural dynamics of GB1 (WT) and ProFORGE-designed proteins at 310 K.** (a–c) PyMOL cartoon snapshots of GB1 (WT), 98g3-0203, and 98g3-0352, respectively, at the predetermined 0-, 100-, 200-, 300-, 400-, and 500-ns checkpoints. (d–f) Residue-by-time maps of three-state secondary structure (SS3) for the same proteins in the same order. (g–i) C$\alpha$ RMSD relative to each condition-specific starting structure after least-squares C$\alpha$ fitting. All three RMSD panels use a common range of 0–0.6 nm. For Supplementary Figs. 5–7, one 500-ns trajectory was analysed for each protein–temperature condition. Frames were sampled at 100-ps intervals, giving 5,001 frames per trajectory. Snapshots were fitted to the corresponding reference structures in main Fig. 5b using all corresponding C$\alpha$ atoms without outlier rejection. Cartoons and SS3 maps use the same GROMACS DSSP assignments, grouped into helix (H), $\beta$-strand (E), and coil/turn (C). These states are shown in pink, ochre, and gray, respectively. Red dashed guides in panels d–i mark 100, 200, 300, 400, and 500 ns.

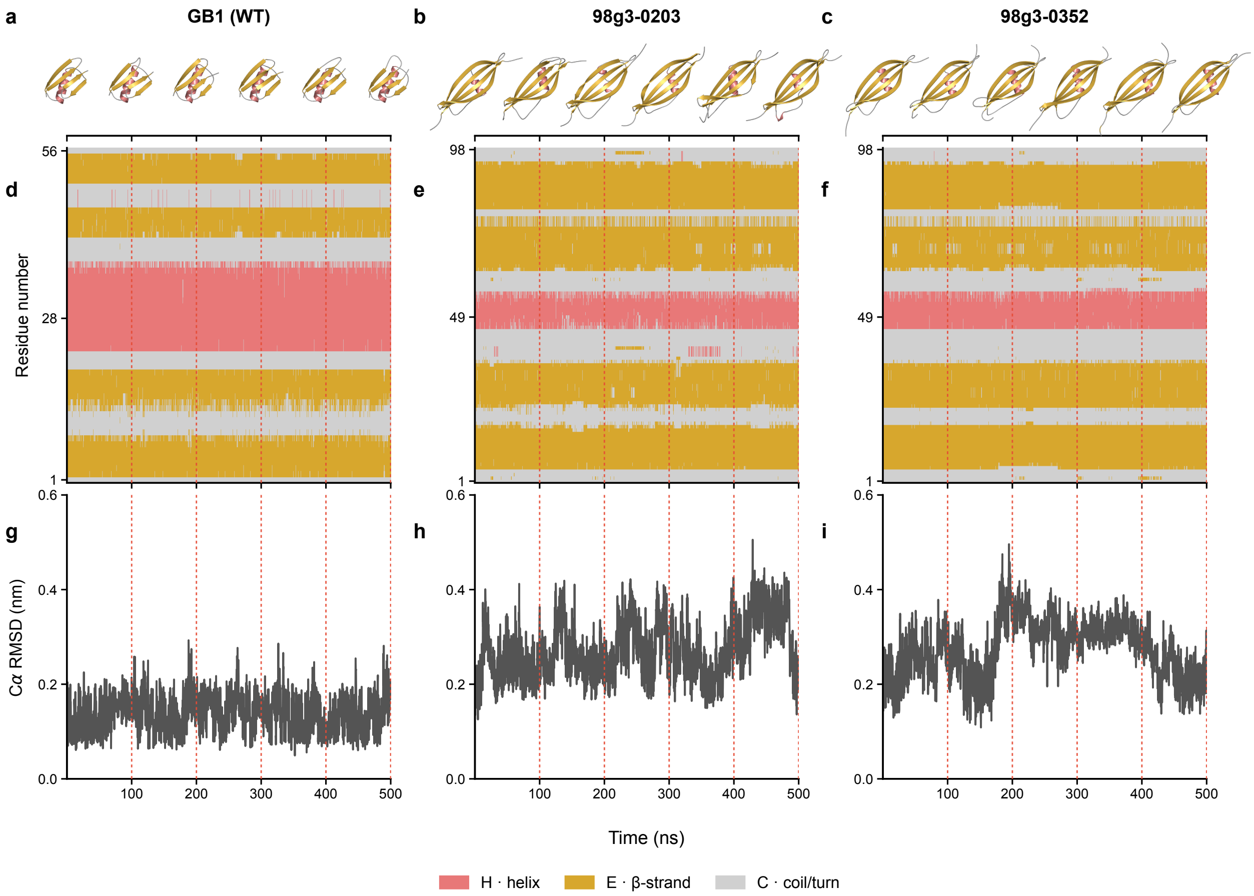


**Supplementary Fig. 6: Structural dynamics of GB1 (WT) and ProFORGE-designed proteins at 373 K.** Panels a–i are arranged and analysed as in Supplementary Fig. 5. All three RMSD panels use a common vertical range of 0–0.6 nm.

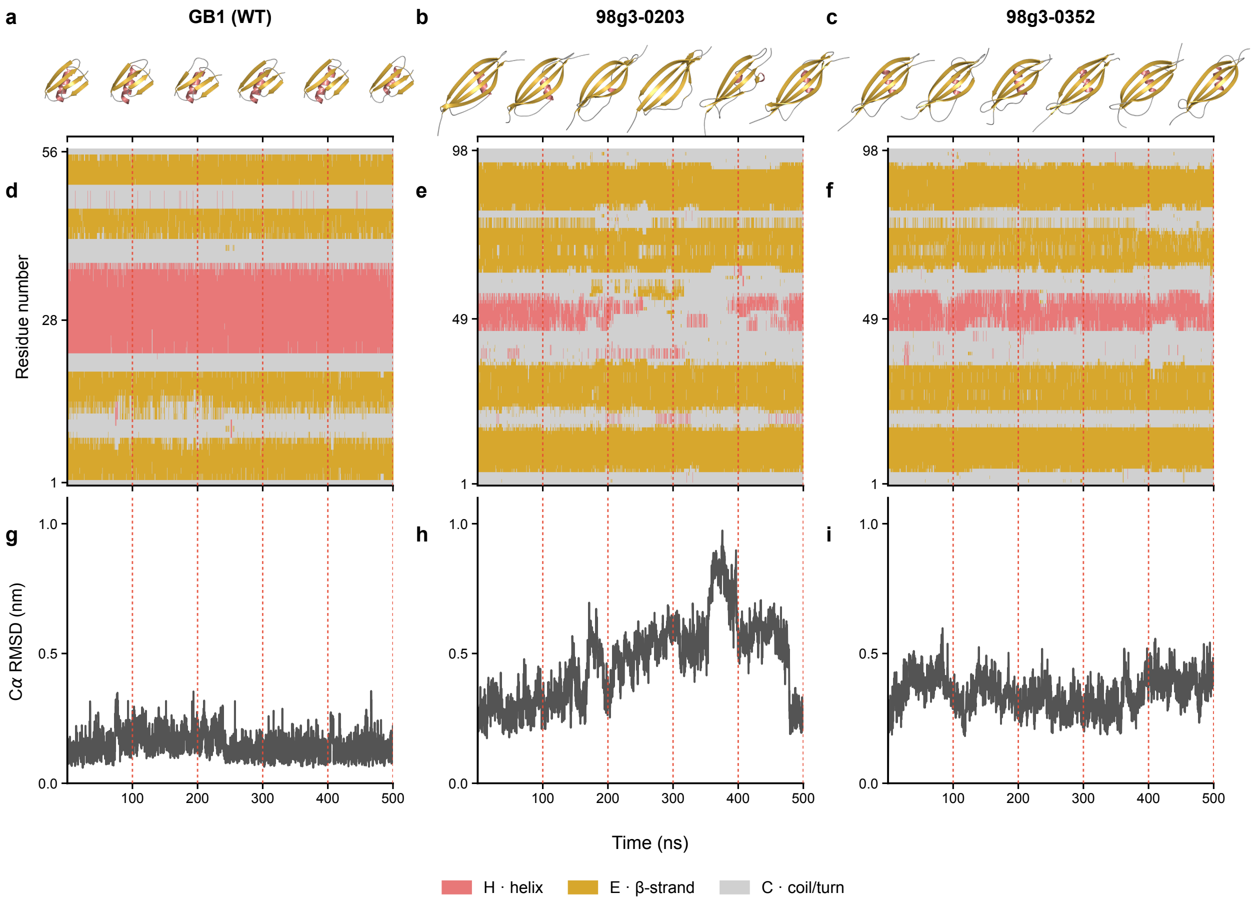


**Supplementary Fig. 7: Structural dynamics of GB1 (WT) and ProFORGE-designed proteins at 430 K.** Panels a–i are arranged and analysed as in Supplementary Fig. 5. All three RMSD panels use a common vertical range of 0–1.1 nm.

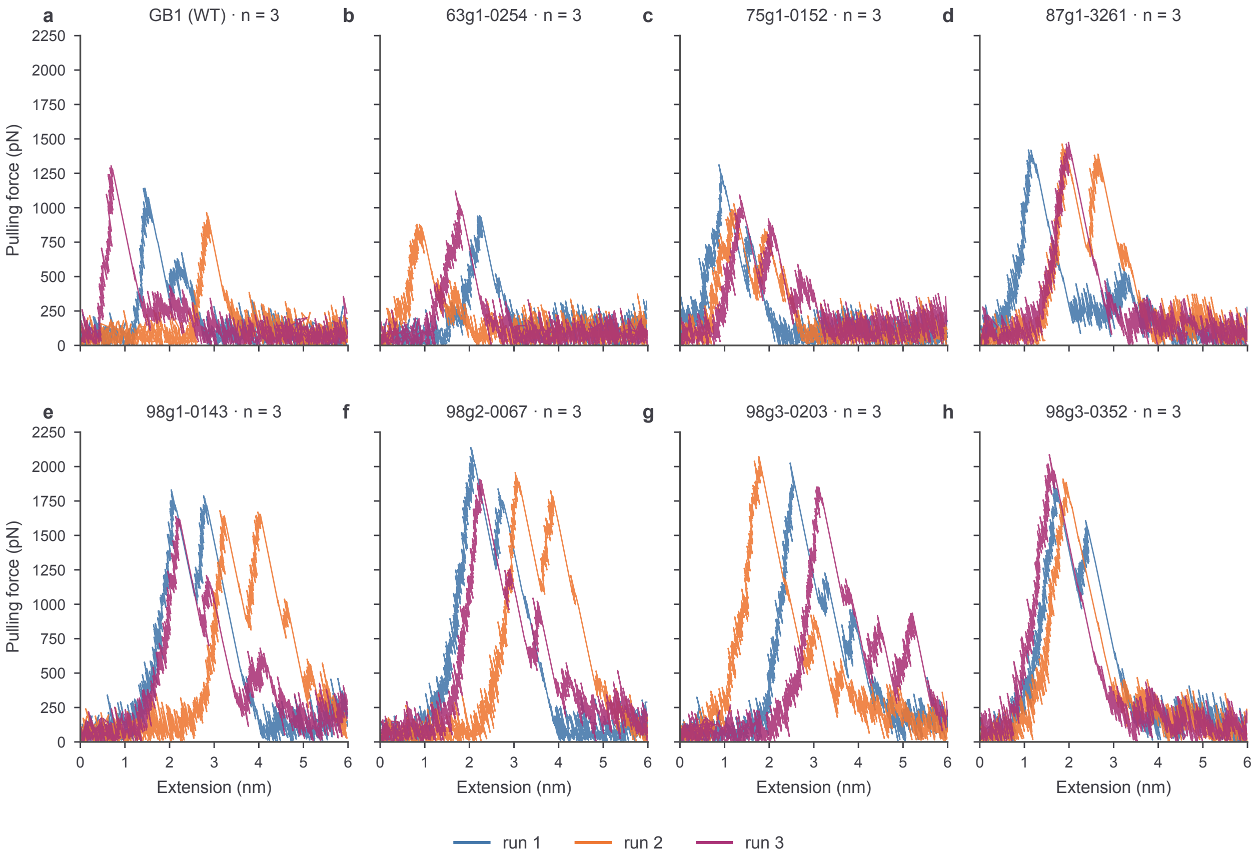


**Supplementary Fig. 8: Raw production-speed SMD force–extension trajectories across the design lineage, with three runs shown per system.** Panels (a)–(h) show GB1 (WT), 63g1-0254, 75g1-0152, 87g1-3261, 98g1-0143, 98g2-0067, 98g3-0203, and 98g3-0352, respectively. The first three runs in numerical run-index order (run 1, run 2, and run 3) are shown in blue, orange, and purple, respectively. Absolute raw pulling force is shown in pN over 0–6 nm extension without temporal smoothing. All displayed trajectories used a pulling speed of 0.5 nm $ns^{-1}$.

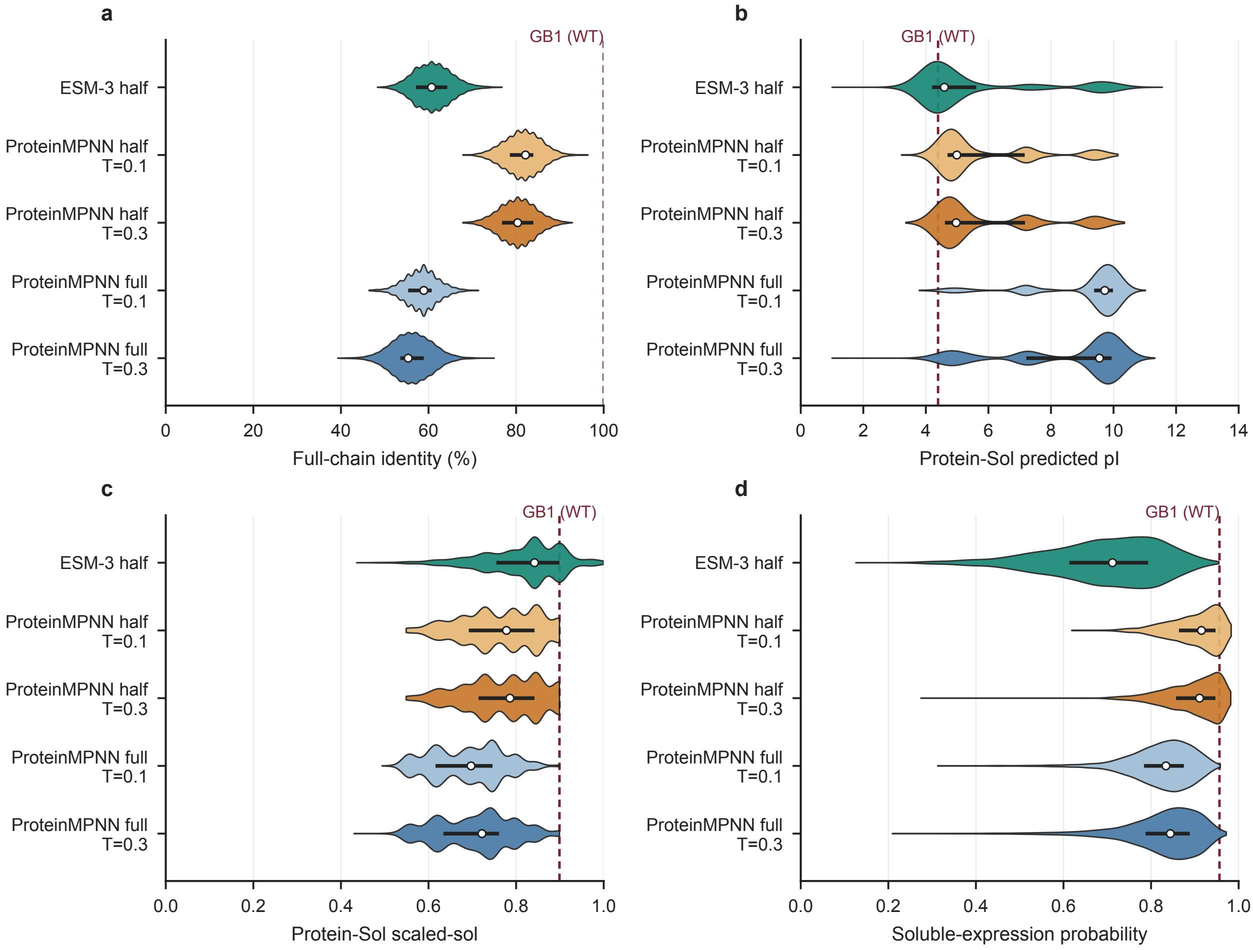


**Supplementary Fig. 9: Sequence-property and predictor-output distributions across five generated-sequence ensembles.** (a) Full-chain identity to GB1 (WT), expressed as a percentage. (b) Protein-Sol-predicted pI. (c) Protein-Sol scaled-sol. (d) SoluProt soluble-expression probability. Horizontal violins show the unique-sequence distributions; white points mark medians, black horizontal bars mark interquartile ranges, and red dashed lines mark the single GB1 (WT) reference. From top to bottom in each panel, the sample sizes are 2,930 (ESM-3), 2,930 and 2,930 (half-masked ProteinMPNN), and 2,927 and 2,930 (fully masked ProteinMPNN).

a

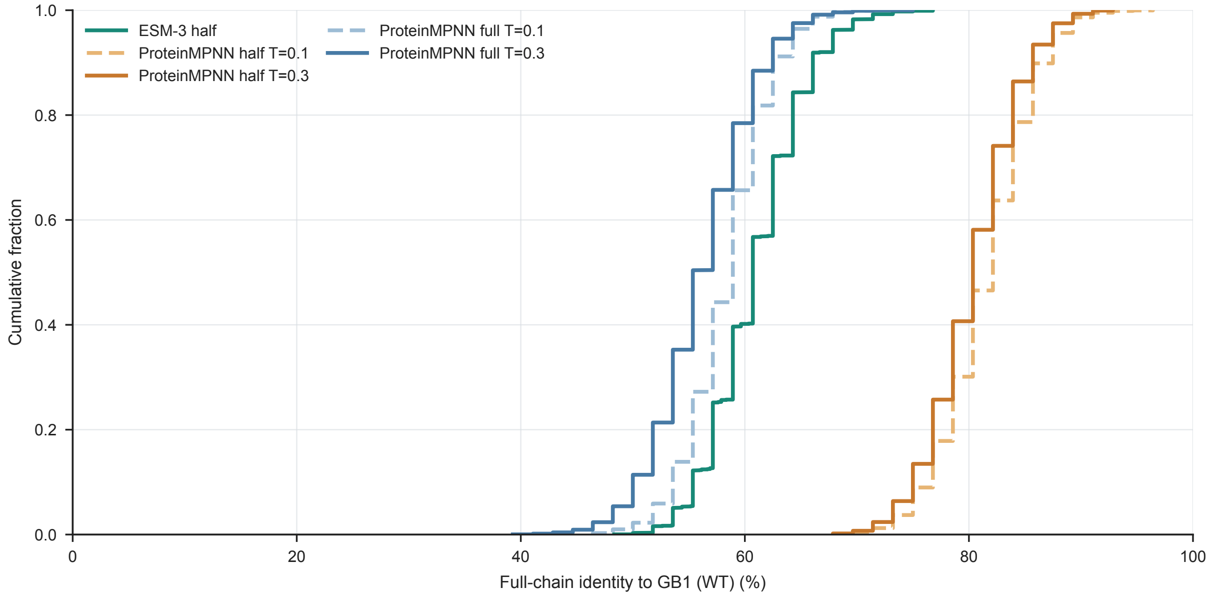


**Supplementary Fig. 10: Empirical full-chain identity distributions across five generated-sequence ensembles.** Panel (a) shows the empirical cumulative distribution of full-chain sequence identity to wild-type GB1 for all unique sequences in each ensemble.

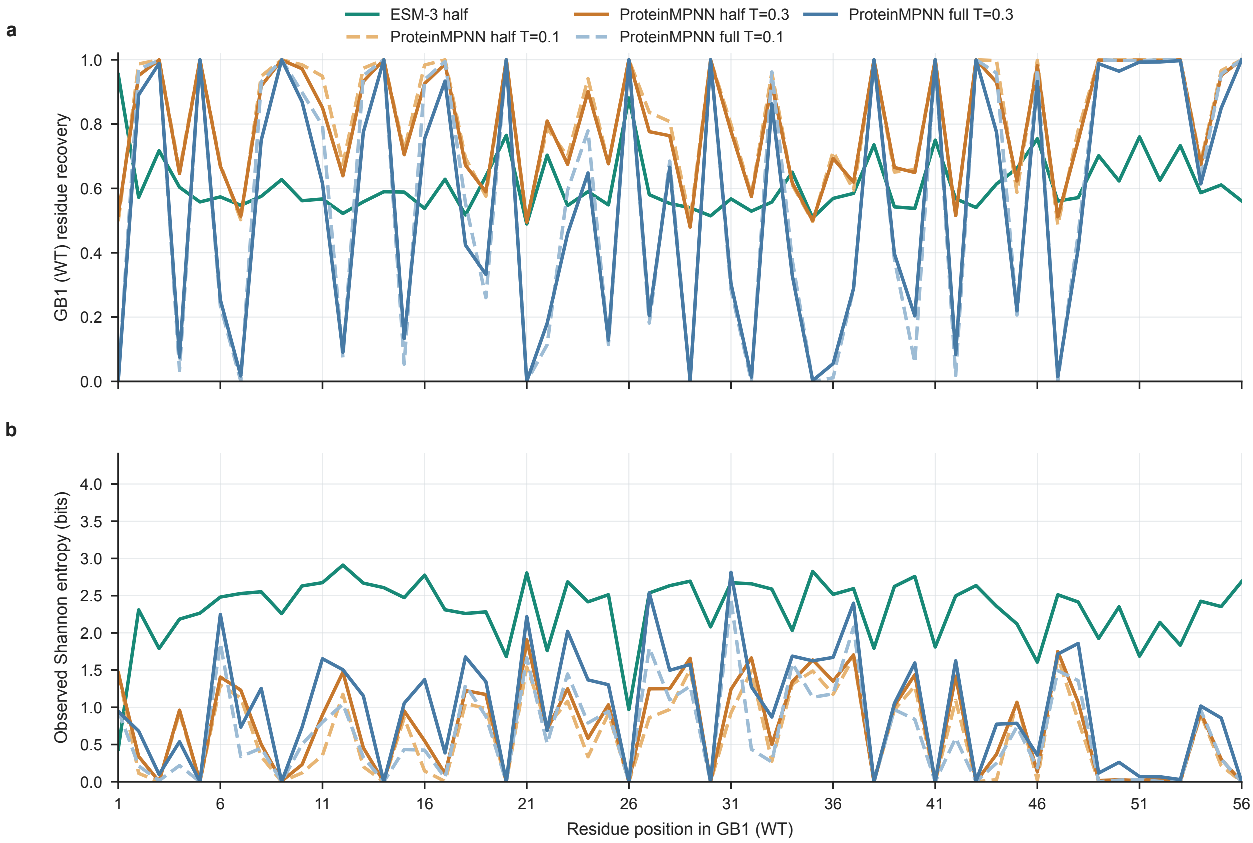


**Supplementary Fig. 11: Position-wise wild-type residue recovery and sequence entropy across five generated-sequence ensembles.** (a) Observed fraction of sequences carrying the GB1 (WT) amino acid at each of the 56 positions. (b) Base-2 Shannon entropy calculated from the raw observed frequencies of the 20 amino acids at each position.

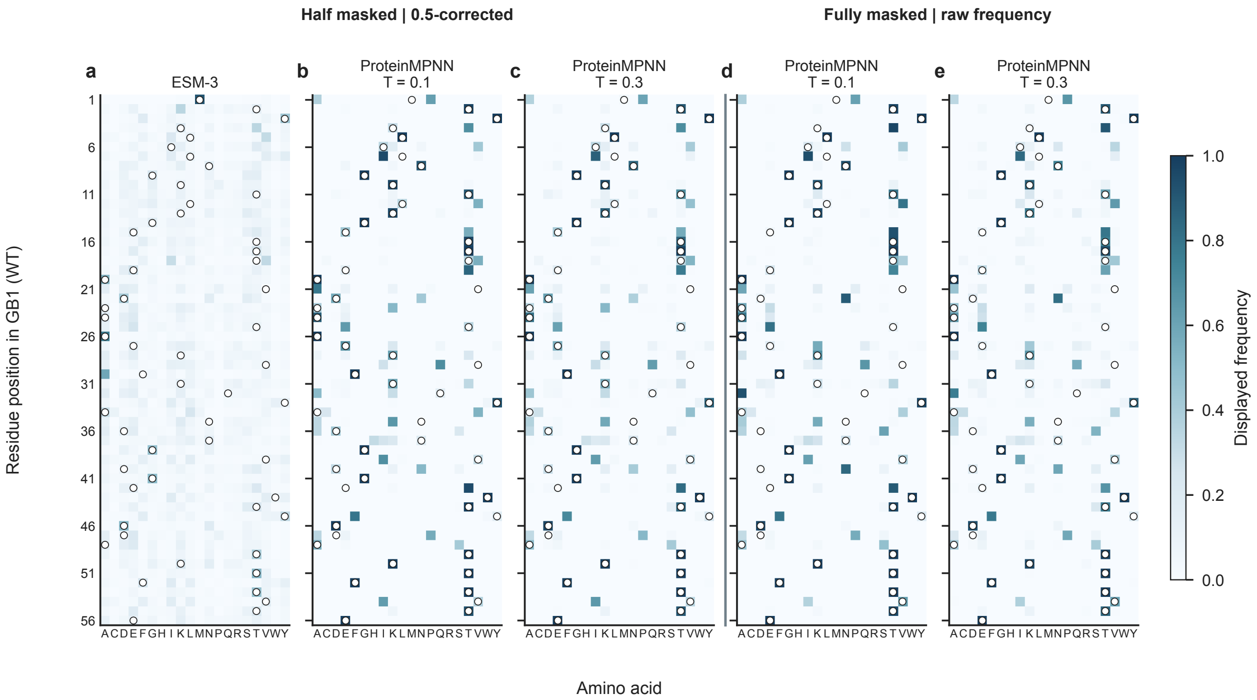


**Supplementary Fig. 12: Adjusted and raw position-by-amino-acid frequencies across five generated-sequence ensembles.** Five heatmaps show the 20 standard amino acids as columns and the 56 GB1 (WT) residue positions as rows. White circles with dark outlines mark the wild-type amino acid at each position. (a) Half-masked ESM-3. (b) Half-masked ProteinMPNN at $T = 0.1$. (c) Half-masked ProteinMPNN at $T = 0.3$. (d) Fully masked ProteinMPNN at $T = 0.1$. (e) Fully masked ProteinMPNN at $T = 0.3$. Panels (a)–(c) show adjusted frequencies: 0.5 was subtracted from the wild-type residue frequency at every position, negative results were set to zero, and the frequencies of the 20 amino acids were renormalized to sum to one at each position. Panels (d)–(e) show raw empirical frequencies, including the wild-type residue frequency at each position. All panels share the same 0–1 displayed-frequency scale.

### Supplementary tables

**Supplementary Table 1: Design lineage and positional constraints.**

| **Parent → cohort** | **Generator** | **Position-control scheme** | **Length (aa)** | **Generated** | **Retained, n/N (%)** | **Redesign, n/N (%)** | **Inserted, n** |
|---|---|---|---|---|---|---|---|
| 1PGA WT → 63g1 cohort | ESM-3 | Batch-shared parent-coordinate redesign and insertion plan | 56 → 63 | 493 | 33/56 (58.93) | 23/56 (41.07) | 7 |
| 63g1-0254 → 75g1 cohort | ESM-3 | Batch-shared parent-coordinate redesign and insertion plan | 63 → 75 | 1,410 | 41/63 (65.08) | 22/63 (34.92) | 12 |
| 75g1-0152 → 87g1 cohort | ESM-3 | Batch-shared parent-coordinate redesign and insertion plan | 75 → 87 | 6,284 | 43/75 (57.33) | 32/75 (42.67) | 12 |
| 87g1-3261 → 98g1 cohort | ESM-3 | Batch-shared parent-coordinate redesign and insertion plan | 87 → 98 | 3,086 | 53/87 (60.92) | 34/87 (39.08) | 11 |
| 98g1-0143 → 98g2 cohort | ESM-3 | Candidate-specific random 17-position redesign plan | 98 → 98 | 269 | 81/98 (82.65) | 17/98 (17.35) | 0 |
| 98g2-0067 → 98g3 cohort | ProteinMPNN ($T = 0.3$) | Full-sequence redesign | 98 → 98 | 3,000 | 0/98 (0.00) | 98/98 (100.00) | 0 |

A. Each denominator is the parent or template length. Redesign counts refer to positions eligible for change.

**Supplementary Table 1: Positional coordinates.** Continued.

| **Parent → cohort** | **Retained parent coordinates** | **Redesign-assigned parent coordinates** | **Insertion anchors in parent coordinates** | **Inserted positions in output** |
|---|---|---|---|---|
| 1PGA WT → 63g1 cohort | 3–4, 6–7, 10–11, 13–15, 17–18, 21, 24–25, 27, 29–31, 33–35, 37–40, 43–45, 47–49, 52, 54 | 1–2, 5, 8–9, 12, 16, 19–20, 22–23, 26, 28, 32, 36, 41–42, 46, 50–51, 53, 55–56 | After parent residues 1, 7, 20, 21, 40, 51 and 56 | 2, 9, 23, 25, 45, 57, 63 |
| 63g1-0254 → 75g1 cohort | 3–15, 18–23, 28, 31–34, 37–39, 42, 45–50, 53–54, 57–61 | 1–2, 16–17, 24–27, 29–30, 35–36, 40–41, 43–44, 51–52, 55–56, 62–63 | One before parent residue 1; one after each of parent residues 16, 25, 27, 29, 36, 41, 42, 51, 54, 61, and 62 | 1, 18, 28, 31, 34, 42, 48, 50, 60, 64, 72, 74 |
| 75g1-0152 → 87g1 cohort | 3–5, 8–10, 13–15, 18–20, 23–24, 27–30, 33–35, 38–41, 44–46, 49, 52–54, 57–58, 61–63, 66–67, 69–72 | 1–2, 6–7, 11–12, 16–17, 21–22, 25–26, 31–32, 36–37, 42–43, 47–48, 50–51, 55–56, 59–60, 64–65, 68, 73–75 | After parent residues 1, 10, 16, 25, 30, 35, 43, 48, 50, 60, 65, and 75 | 2, 12, 19, 29, 35, 41, 50, 56, 59, 70, 76, 87 |
| 87g1-3261 → 98g1 cohort | 4–7, 9, 11–14, 18, 22–26, 28–31, 33–35, 38–40, 42–43, 46–47, 49–52, 55–56, 60–63, 65–67, 69–70, 73–74, 77–79, 81–83, 85 | 1–3, 8, 10, 15–17, 19–21, 27, 32, 36–37, 41, 44–45, 48, 53–54, 57–59, 64, 68, 71–72, 75–76, 80, 84, 86–87 | One after each of parent residues 2, 3, 37, 44, 52, 74, and 86; four after parent residue 87 | 3, 5, 40, 48, 57, 80, 93, 95–98 |
| 98g1-0143 → 98g2 cohort | Complement of the candidate-specific 17-position subset; 81 positions per candidate | Candidate-specific 17-position subset selected from parent positions 1–98 | None | None |
| 98g2-0067 → 98g3 cohort | None | 1–98 | None | None |

B. All positions are one-based.

**Supplementary Table 2: Generation-specific screening criteria.**

| Round | Mean C$\alpha$ pLDDT | Region 1 | Region 2 | Min. E run in each of 1, 2 | Region 3 | Region 4 | Min. E run in each of 3, 4 | Protein-Sol scaled-sol | SoluProt probability |
|---|---|---|---|---|---|---|---|---|---|
| 63g1 | $> 90$ | 2–10 | 56–62 | $\geq$7 | 15–22 | 46–51 | $\geq$6 | $\geq 0.70$ | $\geq 0.70$ |
| 75g1 | $> 90$ | 1–12 | 64–75 | $\geq$9 | 17–28 | 50–61 | $\geq$9 | $\geq 0.80$ | $\geq 0.80$ |
| 87g1 | $> 90$ | 1–14 | 74–87 | $\geq 11$ | 18–30 | 58–70 | $\geq$9 | $\geq 0.75$ | $\geq 0.75$ |
| 98g1 | $> 85$ | 1–21 | 78–98 | $\geq 12$ | 20–39 | 60–79 | $\geq 11$ | $\geq 0.80$ | $\geq 0.80$ |
| 98g2 | $> 85$ | 1–21 | 78–98 | $\geq 12$ | 20–39 | 60–79 | $\geq 11$ | $\geq 0.80$ | $\geq 0.80$ |
| 98g3 | $> 90$ | 1–21 | 78–98 | $\geq 13$ | 20–39 | 60–79 | $\geq 11$ | $\geq 0.80$ | $\geq 0.80$ |

DSSP E runs are contiguous. Each interval is inclusive and must pass independently. Both solubility thresholds must be met.

**Supplementary Table 3: Candidate counts at each screening stage.**

| Assessment | Sequences entering | Sequences retained |
|---|---|---|
| 63g1 combined screen | 493 | 19 |
| 75g1 combined screen | 1,410 | 13 |
| 87g1 combined screen | 6,284 | 23 |
| 98g1 combined screen | 3,086 | 24 |
| 98g2 combined screen | 269 | 66 |
| 98g3 combined screen | 3,000 | 182 |
| 98g3 external AlphaFold 3 assessment | 182 | 19 |

Only the final round underwent the additional AlphaFold 3 screen (mean C$\alpha$ pLDDT $>$ 85).

**Supplementary Table 4: Design-centred sequence and structure comparisons.**

| Pair | Alignment length | Identity, n/N (%) | Similarity, n/N (%) | Gaps, n/N (%) | TM-score | RMSD (Å) |
|---|---|---|---|---|---|---|
| 98g3-0203 vs 98g3-0352 | 98 | 49/98 (50.0) | 66/98 (67.3) | 0/98 (0.0) | 0.9297 | 1.149 |
| 98g3-0203 vs 3GB1 WT | 121 | 12/121 (9.9) | 19/121 (15.7) | 88/121 (72.7) | 0.6376 | 2.472 |
| 98g3-0203 vs B42 | 122 | 23/122 (18.9) | 40/122 (32.8) | 49/122 (40.2) | 0.4960 | 3.434 |
| 98g3-0203 vs B347 | 105 | 24/105 (22.9) | 45/105 (42.9) | 15/105 (14.3) | 0.4965 | 3.396 |
| 98g3-0352 vs 3GB1 WT | 108 | 10/108 (9.3) | 24/108 (22.2) | 62/108 (57.4) | 0.6409 | 2.564 |
| 98g3-0352 vs B42 | 112 | 23/112 (20.5) | 45/112 (40.2) | 29/112 (25.9) | 0.5137 | 3.350 |
| 98g3-0352 vs B347 | 111 | 28/111 (25.2) | 50/111 (45.0) | 27/111 (24.3) | 0.5187 | 2.923 |

EMBOSS Needle percentages use the full global alignment length. TM-score and RMSD are from mTM-align.

**Supplementary Table 5: SMD system construction and equilibration.**

| Stage | Parameter | Fast screen | Detailed and production |
|---|---|---|---|
| System construction | Force field | CHARMM36m | CHARMM36m |
| | Water model | TIP3P | TIP3P |
| | Ionic conditions | Neutralized; 0.15 M NaCl | Neutralized; 0.15 M NaCl |
| | Periodic box | 8 nm $\times$ 8 nm in x and y; z sized for the pulling path and buffer | 8 nm $\times$ 8 nm in x and y; z sized for the pulling path and buffer |
| Energy minimization | Algorithm | Steepest descent | Steepest descent |
| | Maximum steps | 1,000,000 | 1,000,000 |
| | Convergence criterion | Fmax $<$ 1,000 kJ mol$^{-1}$ nm$^{-1}$ | Fmax $<$ 1,000 kJ mol$^{-1}$ nm$^{-1}$ |
| | Position restraints | Backbone 400; side chain 40 kJ mol$^{-1}$ nm$^{-2}$ | Backbone 400; side chain 40 kJ mol$^{-1}$ nm$^{-2}$ |
| NVT equilibration | Duration | 125 ps | 125 ps |
| | Time step | 0.001 ps | 0.001 ps |
| | Temperature coupling | V-rescale; 310 K; $\tau$t = 1 ps | V-rescale; 310 K; $\tau$t = 1 ps |
| | Constraints | Hydrogen bonds with LINCS | Hydrogen bonds with LINCS |
| | Nonbonded treatment | Verlet; 1.2 nm cutoffs; PME electrostatics | Verlet; 1.2 nm cutoffs; PME electrostatics |
| NPT equilibration | Duration | $\geq$ 100 ps | 1 ns |
| | Time step | 0.002 ps | 0.002 ps |
| | Number of steps | $\geq$ 50,000 | 500,000 |
| | Temperature coupling | V-rescale; 310 K; $\tau$t = 1 ps | V-rescale; 310 K; $\tau$t = 1 ps |
| | Pressure coupling | C-rescale; isotropic; 1 bar; $\tau$p = 5 ps; compressibility 4.5 $\times$ 10$^{-5}$ bar$^{-1}$ | C-rescale; isotropic; 1 bar; $\tau$p = 5 ps; compressibility 4.5 $\times$ 10$^{-5}$ bar$^{-1}$ |
| | Constraints | Hydrogen bonds with LINCS | Hydrogen bonds with LINCS |
| | Nonbonded treatment | Verlet; 1.2 nm cutoffs; PME electrostatics | Verlet; 1.2 nm cutoffs; PME electrostatics |

A. Rough-screening NPT equilibration lasted at least 100 ps. Longer runs used more integration steps.

**Supplementary Table 5: Pulling and output sampling.** Continued.

| **Stage** | **Parameter** | **Fast screen** | **Detailed and production** |
|---|---|---|---|
| Steered molecular dynamics | Integrator | md | md |
| | Time step | 0.002 ps | 0.002 ps |
| | Number of steps | 100,000 | 10,000,000 |
| | Duration | 200 ps | 20 ns |
| | Pulling rate | 0.05 nm $ps^{-1}$ (50 nm $ns^{-1}$) | 0.0005 nm $ps^{-1}$ (0.5 nm $ns^{-1}$) |
| | Nominal pulling distance | Approximately 10 nm | Approximately 10 nm |
| | Pull groups | N-terminal C$\alpha$ restrained; C-terminal C$\alpha$ pulled | N-terminal C$\alpha$ restrained; C-terminal C$\alpha$ pulled |
| | Pull geometry | Direction-periodic along +z | Direction-periodic along +z |
| | Spring constant | 1,000 kJ $mol^{-1}$ $nm^{-2}$ | 1,000 kJ $mol^{-1}$ $nm^{-2}$ |
| | Constraints | All bonds with LINCS | All bonds with LINCS |
| | Temperature coupling | Nose–Hoover; 310 K; $\tau$t = 1 ps | Nose–Hoover; 310 K; $\tau$t = 1 ps |
| | Pressure coupling | Parrinello–Rahman; semi-isotropic; 1 bar; $\tau$p = 5 ps; compressibility $4.5 \times 10^{-5}$ $bar^{-1}$ in x/y and 0 along z | Parrinello–Rahman; semi-isotropic; 1 bar; $\tau$p = 5 ps; compressibility $4.5 \times 10^{-5}$ $bar^{-1}$ in x/y and 0 along z |
| | Neighbor searching | Verlet; nstlist=20; rlist = 1.4 nm | Verlet; nstlist=20; rlist = 1.4 nm |
| | Short-range cutoffs | rCoulomb = 1.4 nm; rvdW = 1.4 nm | rCoulomb = 1.4 nm; rvdW = 1.4 nm |
| | Electrostatics | PME; Fourier spacing 0.12 nm; interpolation order 4 | PME; Fourier spacing 0.12 nm; interpolation order 4 |
| Output sampling | Compressed coordinates | 10 or 100 ps, depending on batch | 10 ps |
| | Pull force and displacement | 0.1 or 10 ps, depending on batch | 10 ps |
| | Energies | 1 ps | 1 ps |

B. Detailed and production trajectories shared the same physical pulling parameters. Quantitative force comparisons use matched pulling speeds.

**Supplementary Table 6: Sequences of the seven reported designs.**

| Candidate | Length (aa) | Amino acid sequence (N to C) |
|---|---|---|
| 63g1-0254 | 63 | SISYKVILTNKKTEKGELTTTQEVPDDATVEKVFKEYANSNGVDFTNWTYSDATNTWFCVVKK |
| 75g1-0152 | 75 | MHKSYKVILTNKKTEKQFELTTTQETPQESSAEYTEKVFLNQYANKVGGKVQFTNWTYVEDATDETWFCVVEENE |
| 87g1-3261 | 87 | MPQKSYQAILTVYEKTEKGSFELLTTQKAGPQESDDLEYTADEVFLNFAKANKPGPGARLQFTDCTYEVDDATGTVTWHCVVEIAMP |
| 98g1-0143 | 98 | LANERKSYQIICTVYEVNSKDSTELLTTQKAGPYESDDVYEYTFDETSALNWAKANYDVGPNKKLQFTNCTYYVDDKTGTIETWHVVVEIAIVAKANL |
| 98g2-0067 | 98 | LANERKSYQIICKVYELNSKDSTELLTEQKAGPYESDDVYTYTFDETSALNWAKANYDVGPDKKLKFTNCTYYVDDKTGTIETWHVVTEIAIVAKANL |
| 98g3-0203 | 98 | ASTEIKYRKDILYVYKQSDNDEIELLDSKIFGPFPISDEEEFEFSEEEAIEIAKKNLPISKDEEVVFDKYDSYVDKPSSSVSLTINKYYLLIKKKKKK |
| 98g3-0352 | 98 | DKGKKRYRKDIITFKKLNDDNTIEELMSKEFGPYPISEEKAQKLDIDELIEIAKKELDISEDEEVVFESHSSSVDKPGSDTELITNNYTLIIVKKEKE |

Sequences use the standard one-letter amino acid code. Line breaks do not indicate chain breaks.
